# Shared requirement for MYC upstream super-enhancer region in tissue regeneration and cancer

**DOI:** 10.1101/2022.11.04.515076

**Authors:** Inderpreet Sur, Wenshuo Zhao, Jilin Zhang, Margareta Kling Pilström, Björn Rozell, Anna T. Webb, Huaitao Cheng, Ari Ristimäki, Pekka Katajisto, Martin Enge, Helena Rannikmae, Marc de la Roche, Jussi Taipale

## Abstract

Cancer has been characterized as a wound that does not heal. Malignant cells are morphologically distinct from normal proliferating cells, but have extensive similarities to tissues undergoing wound healing and/or regeneration. The mechanistic basis of this similarity has, however, remained enigmatic. Here we show that the genomic region upstream of Myc, which is required for intestinal tumorigenesis, is also required for intestinal regeneration after radiation damage. The region is also critical for growth of adult intestinal cells in 3D organoid culture, indicating that culture conditions that recapitulate most aspects of normal tissue architecture still reprogram normal cells to proliferate using a mechanism similar to that employed by cancer cells. Our results uncover a genetic link between tissue regeneration and tumorigenesis, and establish that normal tissue renewal and regeneration of tissues after severe damage are mechanistically distinct.

**Summary:** Myc regulatory region links regeneration and cancer

## Main

Out of all genomic regions, the ∽500 kb super-enhancer region on 8q24 that regulates MYC expression carries the largest burden of population-level cancer susceptibility^1,2^; it contains several common alleles that increase risk for multiple major forms of human cancer ^1,3-6^. For example, cancer-associated G-allele of SNP rs6983267 is found in 50% and >85% of people with European and African ancestry, respectively^7,8^. This allele alone increases the risk for colorectal and prostate cancers more than 20%, causing hundreds of thousands of cancer deaths globally per year (https://www.wcrf.org/cancer-trends/global-cancer-data-by-country/). We have previously deleted regulatory sequences syntenic to the super-enhancer region in mice ^9^, and demonstrated that the mice are healthy and fertile in spite of greatly reduced levels of MYC expression in epithelia of the breast, prostate and intestine. The mice, however, are highly resistant to intestinal and mammary tumors. Although deletion of the region did not cause major effect on fitness in our assays, the enhancer elements within it are highly conserved between mammalian species ^10-16^. This suggests that this region would also have a beneficial biological function that would balance the negative impact of the increased cancer predisposition.

In addition to being aberrantly expressed at high levels in majority of human cancers ^17,18^, MYC is also induced by tissue damage^19-21^. MYC has similar functions in both tumorigenesis and tissue repair, driving several biochemical processes including increased ribosomal biogenesis and metabolic flux to support the proliferative state. An acute reduction in MYC level can completely block tumorigenesis^22^ while, both a decrease and an increase in MYC level hinders the process of wound healing ^23,24^. However, it is not known if the same regulatory elements control MYC levels during wound healing and tumorigenesis. Since MYC is one of the main drivers of cell proliferation, we hypothesized that the upstream super-enhancer region would be beneficial under some environmental condition that requires rapid cell proliferation, but which the mice fed well and housed in a relatively clean laboratory environment would not encounter. For example, the super-enhancer region could be important for responses to chronic infections, wounding, or other forms of tissue damage^25,26^.

To test this, we subjected the *Myc*^*Δ2-540/Δ2-540*^ mice to ionizing radiation, which induces a uniform and highly reproducible damage to the intestinal lining ^27,28^. Wild-type mice γ−irradiated for a total dose of 12 Grays rapidly developed intestinal damage and lost 7.0 ± 0.8% of their weight in 3 days (n=19); the weight loss stabilized at that time, and the damage was largely repaired by day 5. By contrast, the *Myc*^*Δ2-540/Δ2-540*^ mice appeared unable to repair the damage, as they continued to lose weight beyond 3 days, losing on average 17% by 5 days (n=10) (**Fig. 1a-b**). Morphological analysis of the small intestines confirmed that the damage was repaired in the wild-type mice but, sustained in the *Myc*^*Δ2-540/Δ2-540*^ mice due to failure to regenerate the intestinal crypts (**Fig. 1c, Extended data Fig. 1**). The wild-type mice displayed robust *Myc* induction at 3 days, and a strong proliferative response at 5 days, whereas at the same time points, in the *Myc*^*Δ2-540/Δ2-540*^ mice *Myc* expression was undetectable and there was almost complete absence of proliferating cells (**Fig. 2a-b, Extended data Fig. 2**). Peak induction of *Myc* transcripts occurred 3 days post irradiation (dpi, **Fig. 2c, Extended data Table 1**). The loss of proliferation was specific to the repair/regenerative stimulus since the cellular architecture and expression profile of cells from the small intestine of the *Myc*^*Δ2-540/Δ2-540*^ mice and wildtype were similar prior to irradiation (**Extended data Fig. 3**). To decipher changes in cellular transcriptome when MYC levels are increasing after tissue damage, we performed single-cell RNA-seq (scRNA-seq) analysis at day 2 after irradiation. Analysis of scRNA-seq data revealed that the *Myc* expression was limited to stem and transit amplifying cells of wild-type mice. Despite the irradiation, which robustly induced MYC in wild-type mice, stem cells or transit amplifying cells from *Myc*^*Δ2-540/Δ2-540*^ mice expressed almost no *Myc*, the signal being even lower than that from irradiated mature enterocytes of wild-type mice (**Fig. 2d**). The *Myc*^*Δ2- 540/Δ2-540*^ mice were, however, fully capable of mounting a normal DNA damage response as seen by the induction of p53 target genes *Cdkn1a* (p21) and *Bbc3* (PUMA), which facilitate p53 dependent cell cycle arrest and apoptosis, respectively (**Fig. 2e**).

**Fig. 1.**
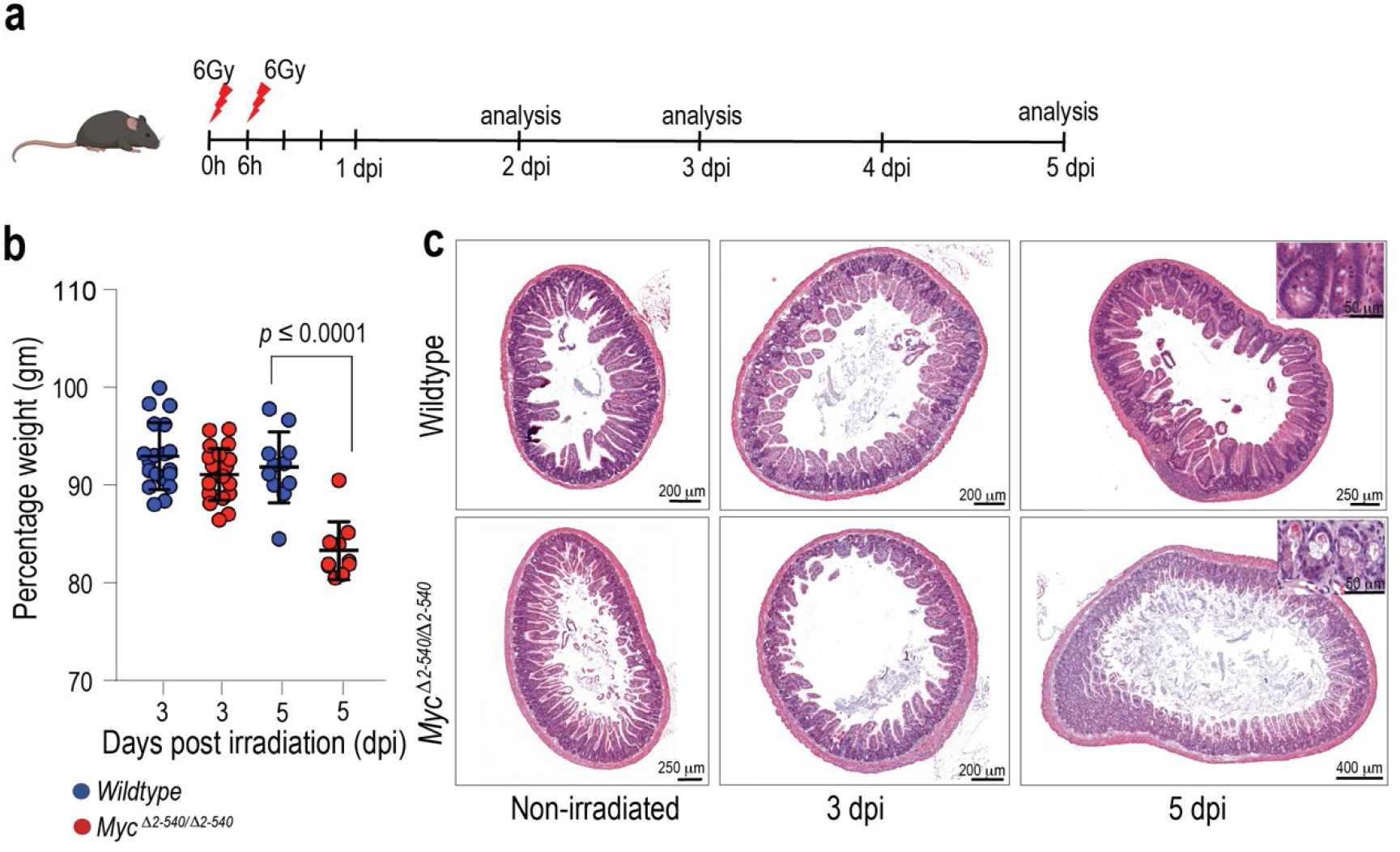
*Myc*^*Δ2-540/Δ2-540*^ intestine fails to support regenerative growth *in vivo* post γ-irradiation. **a**, Timeline of the irradiation experiment. **b**, *Myc*^*Δ2-540/Δ2-540*^ mice (red) are more sensitive to weight loss post irradiation compared to wildtype mice (blue). Wildtype: n = 19 (3dpi), n = 11 (5dpi), *Myc*^*Δ2-540/Δ2-540*^: n = 21 (3dpi), n = 10 (5dpi). Statistical significance was calculated using unpaired student’s *t-test* (two-tailed). **c**, Hematoxylin and Eosin (H&E) stained sections of small intestine showing *Myc*^*Δ2-540/Δ2-540*^ mice fail to regenerate crypts after radiation induced damage.

**Fig. 2.**
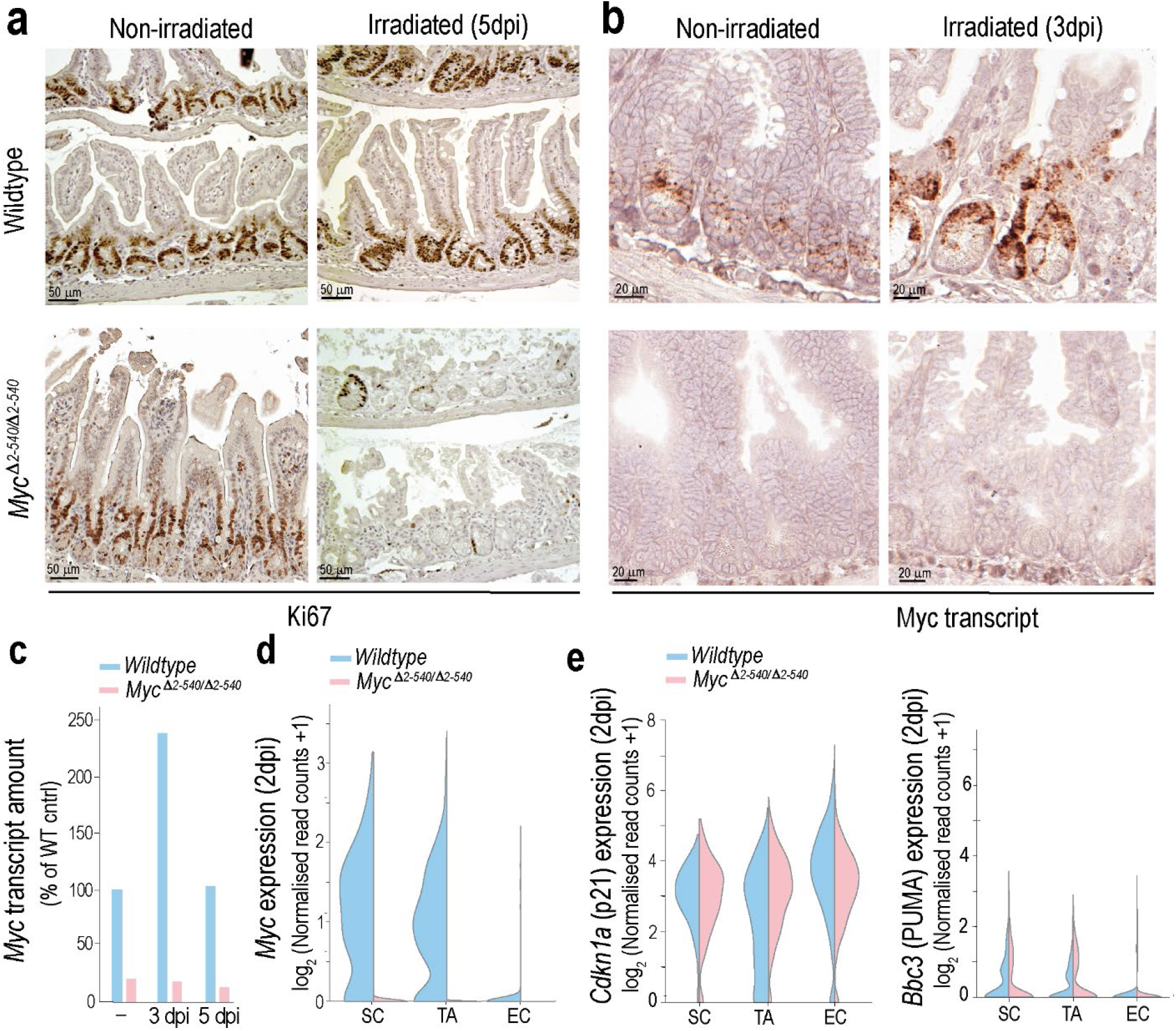
*Myc*^*Δ2-540/Δ2-540*^ intestine fails to induce *Myc* and regenerative proliferation despite a normal p53-mediated DNA damage response post γ-irradiation. **a**, Failure to regenerate the crypts is associated with loss of proliferation as shown by IHC for proliferation marker Ki67. **b-d**, *Myc*^*Δ2-540/Δ2-540*^ intestine is unable to upregulate *Myc* expression after irradiation compared to the wildtype (**b**): ISH; (**c**): qPCR, wildtype: n=2 (control), n=3 (3dpi), n=3 (5dpi); *Myc*^*Δ2-540/Δ2-540*^: n=2 (control), n=2 (3dpi), n=3 (5dpi); (**d**): scRNA-seq, violin plots depicting expression of *Myc* in SC, TA and EC populations are shown. **e**, violin plots showing that unlike *Myc* levels, the levels of p53-mediated DNA damage response indicators *Cdkn1a* (p21) and *Bbc3* (PUMA) were comparable between wildtype and *Myc*^*Δ2-540/Δ2-540*^ intestine at 2dpi (scRNA-seq). SC: stem cell, TA: transit amplifying cell, EC: enterocyte.

**Fig. 3.**
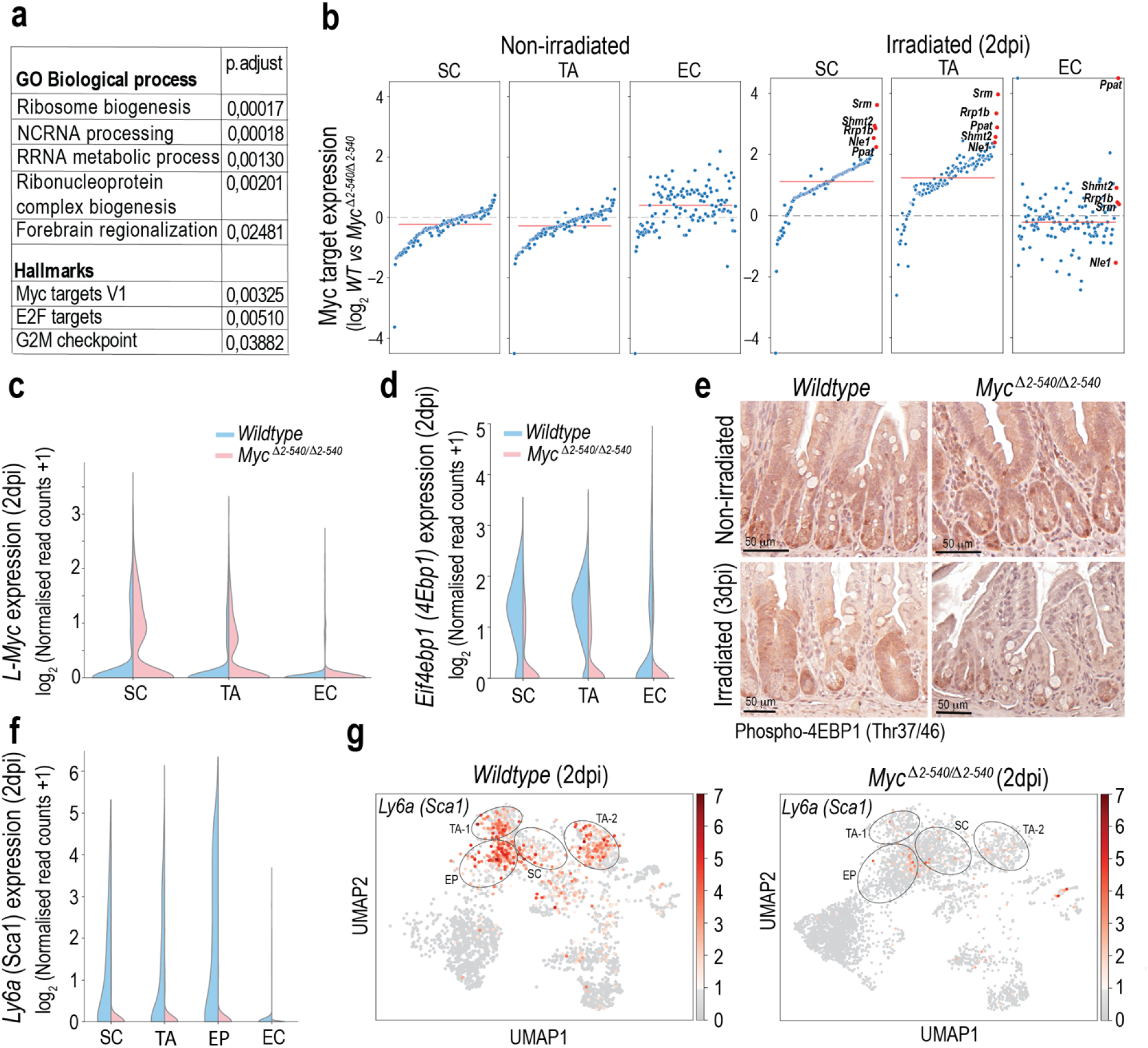
*Myc*^*Δ2-540/Δ2-540*^ intestine fails to induce MYC targets after irradiation, compromising mTORC signalling and stem cell mobilization. **a**, GSEA analysis showing the most enriched gene ontology and hallmarks gene sets in the intestinal SC cluster of irradiated wildtype mice compared to *Myc*^*Δ2-540/Δ2-540*^ mice. **b**, Deletion of Myc^2-540^ region causes a modest decrease in conserved MYC targets expression whereas after irradiation strong induction of MYC targets is seen in wildtype compared to *Myc*^*Δ2-540/Δ2-540*^ mice. Red line denotes the median expression of MYC targets. Red dots denote the top 5 differentially induced MYC targets in the stem cell compartment. **c**, Modest decrease of MYC targets during homeostasis is associated with increased expression of *L-Myc* in the intestinal stem and transit amplifying cells of *Myc*^*Δ2-540/Δ2-540*^ mice compared to wildtypes (also see Extended data Fig. 4). **d-e**, Altered regulation of MYC and mTORC target Eif4ebp1 (4EBP1) (d) scRNA-seq data (e) IHC showiyng amount of phosphorylated 4EBP1 before and after radiation. **f-g**, scRNA-seq data: violin plots showing loss of Ly6a/Sca1 induction in *Myc*^*Δ2-540/Δ2-540*^ mice compared to wildtype after irradiation. SC: stem cell; TA: transit amplifying cell, EP: enterocyte progenitor; EC: enterocyte.

To determine if the loss of regenerative potential was due to MYC, we further analyzed the expression of all genes within 5 Mb of *Myc*. By far, the strongest effect seen was for *Myc* itself, whose expression was almost completely abolished in the mutant intestine. Transcripts of some genes telomeric of *Myc* are also downregulated, but to a much lesser extent. These include the non-coding RNA *Pvt1*, and the gasdermin C genes *Gsdmc2, 3* and *4* (**Extended data Table 2**). However, only the expression of *Myc* and *Pvt1* was specifically upregulated in wildtype cells upon irradiation. Because *Pvt1* lncRNA is thought to respond to the same enhancer elements and elicit its effect via interaction with *Myc* ^29,30^, the results indicate that MYC is the primary effector of the *Myc*^*Δ2-540/Δ2-540*^ phenotype. Consistent with this, unbiased gene set enrichment analysis revealed that a set of MYC targets (V1) failed to be upregulated by irradiation in stem cells of the *Myc*^*Δ2- 540/Δ2-540*^ mice (**Fig. 3a, Extended data Table 3**). We also specifically analyzed expression of a set of 126 functionally conserved MYC targets from Zielke et al. ^31^, most of which are involved in metabolism and ribosome biogenesis. The regulation of this set of genes is conserved between *Drosophila* and humans. The functionally conserved MYC targets were also strongly upregulated in the stem-cells of irradiated wild-type mice compared to *Myc*^*Δ2-540/Δ2-540*^ mice (**Fig. 3b**). These results establish that induction of MYC drives a robust program of increased ribosome biogenesis and anabolic activity that facilitates growth of new tissue to repair the damage caused by the irradiation. However, among the MYC regulated genes, we also observed induction of the negative translational regulator eukaryotic initiation factor 4E binding protein 1 (*Eif4ebp1*) ^32^ (**Fig. 3d**), suggesting that the induction of MYC is necessary but not sufficient to induce the regenerative response. 4EBP1 is inactivated by phosphorylation by the mTOR complex (mTORC) kinase ^33^, and it has been previously shown that mTORC signalling is required for regeneration of intestine after radiation damage ^34^. Staining of the intestines before radiation damage revealed strong signal for phosphorylated 4EBP1 (Thr37/46) while after radiation staining was more prominent in the regenerating crypts of the wildtype (**Fig. 3e**), consistent with a role for both MYC and mTORC in the regenerative response.

Analysis of cell types and proliferation prior to damage revealed no major differences between wild-type and *Myc*^*Δ2-540/Δ2-540*^ mice (**Extended data Fig. 3**). Furthermore, in the normal non-irradiated *Myc*^*Δ2-540/Δ2-540*^ mice, the functionally conserved MYC targets were only modestly downregulated within the stem and transit amplifying cells compared to wildtype mice (**Fig. 3b, Extended data Table 4**). By contrast, there was no downregulation of the MYC target genes in mature enterocytes, which do not express MYC. We observed an increase in *L-Myc* expression within the stem and transit amplifying cell clusters of *Myc*^*Δ2-540/Δ2-540*^ mice suggesting compensation of MYC function during homeostasis and explaining why the loss of the Myc 2-540 kb region is well tolerated in mice under normal conditions (**Fig. 3c, Extended data Fig. 4**). It is noteworthy that the functionally conserved MYC target gene set that we find enriched after tissue damage was originally detected from tumor tissues ^31^, suggesting that increased ribosomal biogenesis and metabolic flux drive both cancer and tissue repair. These results indicate that the failure of the *Myc*^*Δ2-540/Δ2-540*^ is most likely due to failure to respond to damage, as opposed to differences in tissue structure, morphology, or gene activity prior to the damage.

**Fig. 4.**
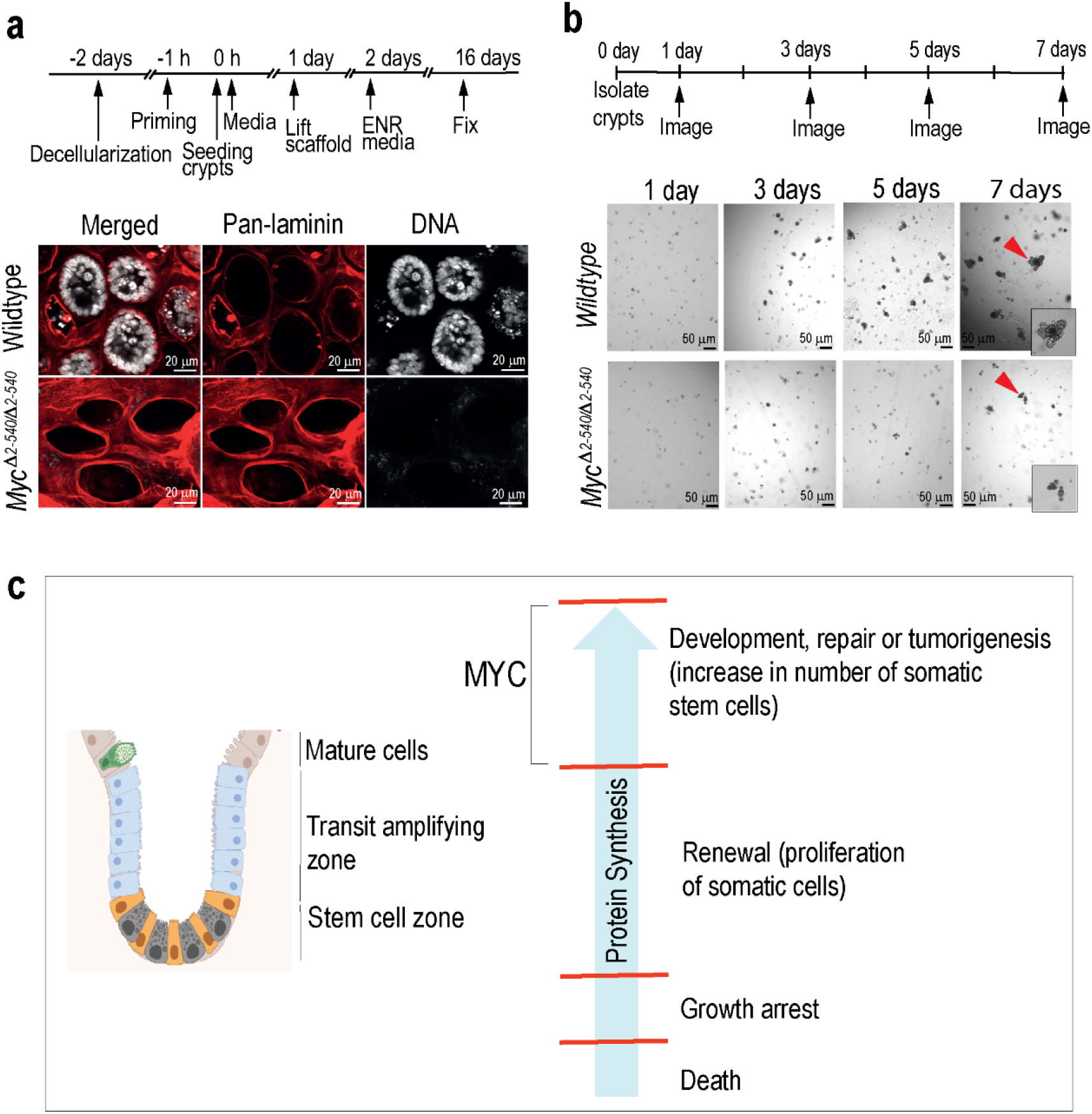
*Myc*^*Δ2-540/Δ2-540*^ intestine fails to support regenerative growth *in vitro* 3-D cultures. **a**, Immunofluorescence analysis showing failure of *Myc*^*Δ2-540/Δ2-540*^ crypts to re-epithelialize intestinal scaffolds. Representative images from two independent experiments are shown. **b**, The inability to re-epithelialize the scaffolds is correlated with the inability of crypts from adult *Myc*^*Δ2-540/Δ2-540*^ intestine to support organoid growth in culture. Representative images from 4 independent experiment are shown. **c**, Model describing how different levels of protein synthesis can result in different proliferative cell fate choices ranging from normal homeostasis to regeneration and cancer.

To further study the mechanism underpinning the failure of *Myc*^*Δ2-540/Δ2-540*^ intestines to regenerate, we determined whether the loss of MYC affected the mobilization of reserve stem/progenitor cells which previously have been shown to be associated with intestinal regeneration ^35,36^. For this purpose, we checked expression of *Ly6a* which codes for the stem cell marker Sca1. Expression of Sca1 is induced by colitis and radiation induced damage as well as damage caused by helminth infections ^35,36^. Analysis of our scRNA-seq data showed that irradiation of the wildtype intestine mobilized a robust population of Ly6a/Sca1 expressing cells. These cells were distributed amongst the stem, transit amplifying and enterocyte progenitor cell clusters, confirming their stem cell potential (**Fig. 3f-g**). By contrast, the *Myc*^*Δ2-540/Δ2-540*^ mice almost completely failed to mobilize the Ly6a/Sca1^+^ cells after irradiation (**Fig. 3f-g**). These results establish that the 2-540 kb conserved region which is required for tumorigenesis ^37^ is also required for MYC-dependent repair of intestinal lining after acute large-scale damage.

To further characterize the role of MYC in tissue repair, we decided to test a recently described *ex vivo* tissue regeneration assay, where mouse ileum is decellularized with sodium deoxycholate resulting in a matrix reminiscent of the 3-D intestinal structure *in vivo* with distinct regions where former crypts and villi were located. Overlaying of the decellularized extracellular matrix (dECM) with isolated crypts results in repopulation of the empty crypt pits and regeneration of the entire intestinal epithelium ^38^. In this assay, cultures of wild-type crypts repopulated the dECM’s and gave rise to crypt and villus structures, whereas crypts from *Myc*^*Δ2-540/Δ2-540*^ mice failed to do so (**Fig. 4a**). As repopulation of the bare dECM requires cellular motility, with proliferation and self-renewal restricted to the empty crypt pits, we next placed isolated crypts in standard organoid culture conditions in matrigel. Surprisingly, adult intestinal organoids from *Myc*^*Δ2-540/Δ2- 540*^ mice failed to grow also in such culture, where self-renewal and proliferation promoting signals are provided evenly (**Fig. 4b**).

Taken together, these results establish that although the *Myc* upstream super-enhancer region is not required for normal tissue renewal in mouse intestine, it is required for tissue repair after radiation damage and for growth of adult intestinal cells under organoid culture conditions.

## Discussion

We show here that the same genetic element, the *Myc* upstream super-enhancer region, is required for both tissue renewal after radiation damage, and tumorigenesis. The super-enhancer region is critical for the induction of *Myc* after intestinal damage caused by irradiation. Mice deficient of the region fail to induce expression of MYC target genes important for tissue repair, and are unable to mobilize Ly6a/Sca1^+^ regenerative stem/progenitor cells. These results establish the long-sought mechanistic link between wound healing and cancer, and show that proliferation during normal homeostasis and tissue regeneration use regulatory mechanisms that are genetically distinct. Given that upregulation of ribosome biogenesis is required for MYC-driven proliferation ^31,39,40^, the simplest model consistent with our findings is that different levels of protein synthesis are needed for distinct proliferative cell fate choices. In this model, relatively low levels of protein synthesis and MYC are sufficient for proliferation that maintains stem cell number in tissue renewal, whereas processes that require exponential increase in the number of somatic stem cells, such as cell proliferation *in vitro*, regeneration, and tumorigenesis (**Fig. 4c**) require high levels of MYC and protein synthesis. Interestingly, the model has similarities to growth control in unicellular species, where nutrient availability specifies the protein synthetic rate and controls the balance between non-proliferative and proliferative states ^41^. Multicellularity could have evolved simply by the addition of a hierarchy of stem cells with new proliferative fates, which remain controlled by the same input variable, the total metabolic activity per genome. Consistently with this model, ES cells lacking MYC enter into a diapause-like non-proliferative state *in vitro* ^42^, and we also observe here that adult intestinal cells from *Myc*^*Δ2-540/Δ2-540*^ mice enter a non-proliferative state under full mitogen stimulation (organoid culture).

Our results also show that cell culture models that are often considered to resemble normal *in vivo* tissue renewal actually drive cells to adopt an exponential cell proliferation program associated with regeneration and tumorigenesis. This has important consequences for drug development. In particular, compounds that inhibit proliferation of normal cells in culture have commonly been discarded during the development process due to toxicity, even when a subset of them may actually target the very mechanism that is distinct between normal proliferative cells and cancer cells.

## Methods

### Animals

Mouse lines used in this study have been described previously ^9^. The conditional knockout mice were generated on a C57BL/6N genetic background. After Cre-mediated deletion of 538 kb region, mice were backcrossed to C57Bl6/J for two generations and subsequently intercrossed. All mice were kept on a 12:12 h light cycle in standard housing conditions and provided with food (Teklad global diet 2918; Envigo) and water *ad libitum*. Animal experiments were done in accordance with the regulations and with approval of the local ethical committee (Stockholms djurförsöksetiska nämnd). Mice were irradiated with Gammacell® 40 Exactor that uses Caesium-137 as a source for γ-rays. Mice were given 2 doses of 6 Gy with a 6 hr interval in between the doses. After irradiation mice were provided with soft food. For measuring survival, the general health of the animal was monitored and mice were sacrificed when they reached the humane end-point according to the ethical permit.

### Immunohistochemistry (IHC) and RNA *In Situ* hybridization (ISH)

PFA-fixed and paraffin embedded (FFPE) tissues were sectioned at 5 μm thickness and subjected to IHC using standard methods. For antigen retrieval, the sections were boiled in microwave for 20 minutes in 10 mM Citrate buffer pH 6.0 and subsequently processed for respective antibody stainings. Following antibodies and working dilutions were used: anti-Ki67 (Abcam, 1:200), anti-Lysozyme (Abcam, 1:500), anti-Muc2 (Abcam, 1:500), anti-Chga (Abcam, 1:1000), anti-phospho 4EBP1 (Cell signalling, 1:2500). ISH was also performed on 5 μm thick sections using the RNAscope technology and according to manufacturer’s protocol. Probes against mouse *Myc* (catalog nr: 413451), *L-Myc* (catalog nr: 552711) and *Lgr5* (catalog nr: 312171) were used. RNA integrity was confirmed using the control probe for *Ubc* (catalog nr:310771).

### qPCR

Approximately 1 cm of the mouse distal ileum was cut into small pieces and stored in RNAlater (Qiagen). RNA was prepared using a combination of Trizol (Ambion) and RNeasy mini kit (Qiagen). cDNA was synthesized from either 1 μg (tissue) or 100 ng (organoid) of total RNA using High Capacity cDNA Reverse Transcription Kit (Applied Biosystems). qPCR was performed using SYBR™ Green master mix (Thermo Fisher) on a LightCycler® 480 instrument (Roche). Primers for *Myc* and *Actin* were as described previously ^9^.

### Single-cell RNA-seq (scRNA-seq)

Mouse intestinal crypts were isolated according to Sato and Clevers ^43^, dissociated with TrypLE express (37°C/20 min, Invitrogen) and single cell suspensions were run on a 10x Chromium controller using the Chromium Next GEM Single Cell 3′kit v3.1 (catalog nr: PN-1000268) and according to manufacturer’s protocol (10x Genomics). Cell number targeted was 6000. For non-irradiated samples, crypts were isolated from ileum and single cell suspension of FACS sorted 7AAD^−^ (Life Technologies), CD45^−^ (eBioscience), CD31^−^ (eBioscience), TER-119^−^ (eBioscience), EpCAM^+^ (eBioscience) cells were used. For irradiated samples, crypts from the whole small intestine were isolated and 7AAD^−^ FACS sorted cells were used. Libraries were sequenced on illumina sequencing platform NovaSeq 6000 or NextSeq 2000. Sequencing data was processed using Cell Ranger (v6.0.1) (10x Genomics). Quality control (QC) analysis of the Cell Ranger output was done in Scanpy (v1.9.1) ^44^ as follows: Mean and Standard deviation (SD) of genes in all cells [Mean (n_genes_), SD(n_genes_)] and Mean, Median, Median absolute deviation (MAD) and SD of total counts in all cells was calculated. Cells were kept with the number of expressed genes larger than [Median(n_genes_)-MAD(n_genes_)], total read counts in the range of [Mean(n_counts_)-1.5*MAD(n_counts_), Mean(n_counts_)+3*SD(n_counts_) and percentage of expressed mitochondrial genes larger than its median+3*MAD. Additionally, genes expressed in less than 0.1% of cells were filtered out. Subsequently for dimensional reduction, top 2000 highly variable genes were selected based on the single-cell RNA raw read count matrix (scanpy.pp.highly_variable_genes; flavor= ‘seurat_v3’). There were no mitochondrial genes in the top 2000 highly variable genes. The raw read count matrix with top 2000 highly variable genes (22,108 × 2,000) was subjected to the Bayesian variational inference model scVI (v0.16.4) ^45^ to infer a latent space for all single cells. The batch effect among different samples was automatically removed for all samples within the scVI. After that, two-dimensional visualizations were calculated by using Uniform Manifold Approximation and Projection (UMAP). Clustering analysis was performed using Leiden algorithm. First a k-nearest-neighbor graph from the scVI latent space was constructed and then sc.tl.leiden function in scanpy was used with the resolution of 1. This resulted in 19 clusters. The small cluster expressing immunocyte markers was not used for further analysis (excluded 34 cells and 22,074 cells were kept). Remaining clusters were annotated into 7 intestinal cell types based on previously published marker genes ^46^.

For further analysis, the single cell raw read count matrix of all the genes was input into R package Scran (v1.24.1) to calculate the size factor for each cell ^47^. To normalize the gene expression across cells, each gene’s read count in a cell was divided by that cell’s size factor in Scanpy. The normalized gene read counts were log_2_ transformed, where 1 pseudo count was added to gene’s read count to avoid negative infinity value in the analysis (log1p value). The z-score for the log1p value for each gene was calculated and used for heatmap visualization of cell-type specific marker genes.

Differentially expressed genes between the genotypes were determined using scanpy’s function sc.tl.rank_genes_groups. The analysis was done specifically in the stem cell (SC), transit amplifying cell (TA) and enterocyte clusters (EC). Within this function, Wilcoxon test was used to calculate p values and the false discovery rate (FDR) by applying the benjamini-hochberg approach. The log fold changes of the 126 known MYC targets ^31^ were retrieved from the differential gene expression results and plotted using a dot plot. Polr1f was not detected in our dataset and was excluded from analysis (**Extended data Table 4**). The ranking of the genes on the X-axis was based on the median log fold change values of each MYC target gene among stem cell (SC), transit amplifying cell (TA) and enterocyte (EC) clusters combined.

Gene Set Enrichment Analysis was done using R package clusterProfiler (v4.4.4). Gene signatures were obtained from Gene Ontology Biological Process gene sets ^48^ and the MSigDB ‘Hallmark gene sets’ ^49^. All the expressed genes in the SC cluster were pre-ranked by their log fold changes which were determined as described in the previous section. GSEA analysis was then performed using ClusterProfiler (ClusterProfiler::GSEA; pvalueCutoff=1, pAdjustMethod=′BH′). The full list of GSEA results is provided in Extended data Table 2.

### Single cell ATAC-seq (scATAC-seq)

scATAC-seq was performed according to Cheng *et al 2021* ^50^. Single cell suspensions of mouse intestinal crypts were prepared as in ^51^ and single cells were FACS sorted into 384 well plate with 3 μl lysis buffer in each well (0.03 μl 1M Tris-pH7.4, 0.0078 μl 5M NaCl, 0.075 μl 10% IGEPAL, 0.075 μl RNases Inhibitor, 0.075 μl 1:1.2M ERCC, 2.7372 μl H_2_O). After cell lysis, 1μl of the lysis buffer containing the nuclei was used for scATAC library preparation, while the remaining 2 μl were discarded. The scATAC *in situ* tagmentation was performed with 2μl of the Tn5 tagmentation mix (0.06 μl 1M Tris-pH 8.0, 0.0405 μl 1M MgCl_2_, 0.2 μl Tn5) at 37°C for 30 mins. After tagmentation, 2 μl of the supernatant was aspirated and the nucleus was then washed once with 10 μl ice-cold washing buffer (0.1 μl 1M Tris-pH7.4, 0.02 μl 5M NaCl, 0.03μl 1M MgCl_2_, 9.85 μl H_2_O). The remaining Tn5 was inactivated by adding 2 μl 0.2% SDS-Triton X-100 and incubating at room temperature for 15 mins followed by 55°C for 10 mins. Before barcoding PCR, genomic DNA was extracted from chromatin by adding 2μl Proteinase K (0.0107 au/mL) and incubation at 55°C for 1h followed by heat inactivation at 70°C for 15 mins. Barcoding PCR was done using KAPA HiFi PCR Kit (Roche) in a final volume of 25 μl. The PCR condition was 72°C/15 min, 95°C/45 s, [98°C/15 s, 67°C/30 s, 72°C/1 min] x 22 cycles, 72°C/5 min, and then 4°C hold. After the PCR, 2μl reaction from each well was pooled and cleaned-up twice using SPRI-beads (at 1.3X volume) and the library was sequenced on Illumina NextSeq 550 as paired-end, dual index (37+37+8+8). For scATAC-seq data, raw sequence reads were trimmed based on quality (phred-scaled value of >20) and the presence of illumina adapters, and then aligned to the mm10 genome build using BWA ^52^. Reads that were not mapped, not with primary alignment, missing a mate, mapq <10, or overlapping ENCODEs blacklist regions ^53^ were removed. A custom script was used to summarize the paired-end reads into a de-duplicated fragment file suitable for downstream analysis. scATAC-seq data was analyzed using ArchR. Cells with fewer than 1000 ATAC-seq reads mapping to transcription start site, or with TSS enrichment < 4, were excluded from analysis. The “iterative LSI” function with maxClusters=6 was used for cluster analysis. Clusters with high expression of ‘Muc2’ were annotated as goblet cells and discarded. Clusters high in ‘Ascl2’ and ‘Lgr5’ were annotated as stem cells, while remaining intermediate/low expressing clusters were annotated as Transit amplifying cells. Each of these superclusters were split by genotype and merged to in-silico bulk experiments.

### Intestinal crypt isolation and Organoid culture

Crypts were isolated from adult mouse small intestine (approx. 3 month old) according to Sato and Clevers 2013 ^43^. The crypts were seeded in Matrigel (Corning) and cultured in ENR medium (Advanced DMEM/F12 (Thermo Fisher Scientific), 1x penicillin/streptomycin, (Thermo Fisher Scientific), 1x Glutamax (Thermo Fisher Scientific), 10mM HEPES (Thermo Fisher Scientific), 1x B27 and 1x N2 (Thermo Fisher Scientific), 1.25mM N-acetylcysteine (Sigma), 50 ng/ml of murine recombinant Epidermal growth factor (R&D), 100 ng/ml recombinant murine Noggin (R&D), 1 mg/ml recombinant murine R-Spondin (R&D).

### Re-epithelialization of intestinal scaffolds with crypts

The ileum from the small intestine of mice, was decellularized as previously described by Iqbal *et al* 2021 ^38^. Pieces of decellularized intestines (dECMs) were primed at 37°C with 50 μl of media containing Advanced DMEM/F12, 1x penicillin/streptomycin, 1x Glutamax (Thermo Fisher Scientific), 10mM HEPES (Thermo Fisher Scientific), 1x B27 (Life Technologies), 1x N2 (Life Technologies), 1mM N-acetylcysteine (Sigma), 50 ng/ml of murine recombinant Epidermal growth factor (R&D), 100 ng/ml recombinant murine Noggin (Peprotech), 500 ng/ml recombinant murine R-Spondin (R&D), 10μM Y-27632 (Sigma), for 15-60 mins. This was followed by aspirating media carefully and plating around 100 crypts on each scaffold piece and incubating at 37°C for 10 min. Next, 10 μl of media was added to each scaffold piece and incubated at 37°C for 10 min, followed by an additional 20 μl per scaffold piece and incubation at 37°C for 20 min. Finally, 260 μl of media was added to each scaffold piece and put in the incubator at 37°C. The day after seeding, media was changed, and each scaffold piece was lifted to make it float. The second day after seeding, the media was changed to media without Y-27632. Media was subsequently replaced every other day. Re-epithelialized dECMs were fixed with 4% PFA at day 16 after plating. The re-epithelialized dECMs were analyzed using immunofluorescence. Primary antibody Pan-laminin (1:300, Abcam, 11575) and secondary antibody Alexa-647 conjugated anti-rabbit (1:1000, Life) were used. For nuclei detection, DAPI (Life, 1:1000) was used. Images were obtained with a Zeiss LSM980-Airy2 microscope using a 20x objective, NA=08.

## Statistics and reproducibility

Sample size was determined based on our previous experience and pilot experiments. Information on statistical significance is provided in the figure legend.

## Data availability

Sequencing data will be uploaded to European Nucleotide Archive (ENA, EMBL-EBI) under project accession number PRJEB57218.

## Acknowledgements

The authors wish to thank Drs Rong Yu, Yujiao Wu and Ekaterina Morgunova for their comments on the manuscript. We also wish to thank the core facilities at KI: The unit for morphological phenotype analysis (FENO), Bioinformatics and expression analysis core facility (BEA), Biomedicum imaging core (BIC) and Biomedicum flow cytometry (BFC). The research was funded by grants from the Swedish Research Council (D0815201) and Cancerfonden (20 1116 PjF 02H).

## Author contribution

IS and JT conceived and designed the experiment, analysed data and wrote the original draft. IKS, MKP, WZ, ATW, HC and HR performed experiments, BR and AR analysed histological data,WZ and JZ performed scRNA-seq data analysis. ME analysed scRNA-seq/scATAC-seq data, PK, IKS, MR, JT provided supervision. WZ, JZ, MKP, BR, AW, HC, ME, AR, PK, HR, MR reviewed and edited the manuscript.

