## Supplemental Figures (1-4) and Tables (1-4) for "Shared requirement for MYC upstream super-enhancer region in tissue regeneration and cancer"

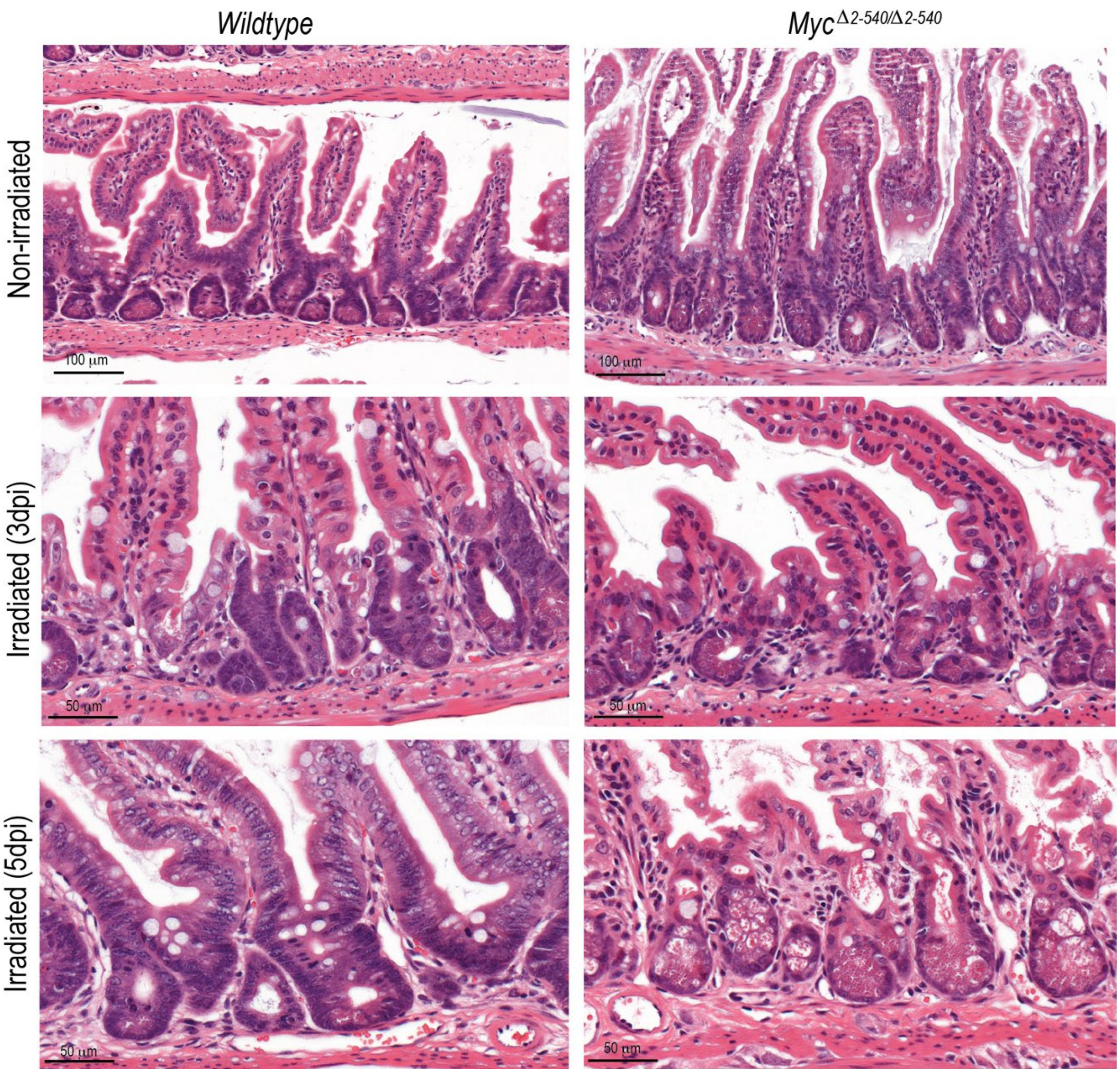


**Extended data Fig. 1**: **Inability of *Myc*$^{\Delta 2\text{-}540/\Delta 2\text{-}540}$ crypts to regenerate after damage caused by γ-irradiation**. Higher magnification images of H&E stained sections as in **Fig. 1c** are shown.

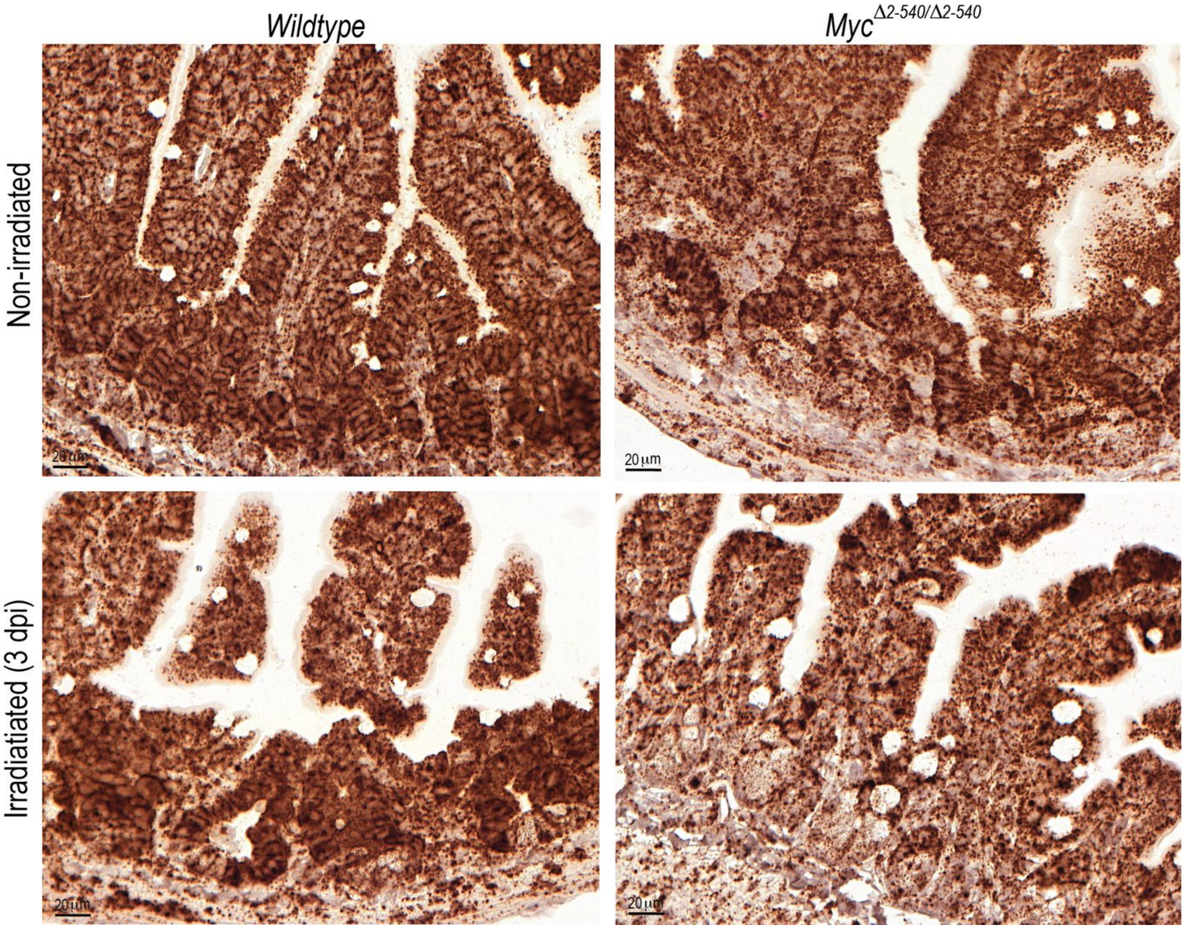


**Extended data Fig. 2**: **Comparable RNA-integrity in ISH sections from wildtype and *Myc*$^{\Delta 2\text{-}540/\Delta 2\text{-}540}$ intestine.** Expression of *Ubiquitin C* (*Ubc*) transcript in the intestine of non-irradiated mice and in mice 3 days after irradiation (3dpi) is shown. *Ubc* transcripts were used as a positive control for RNA integrity of the sample.

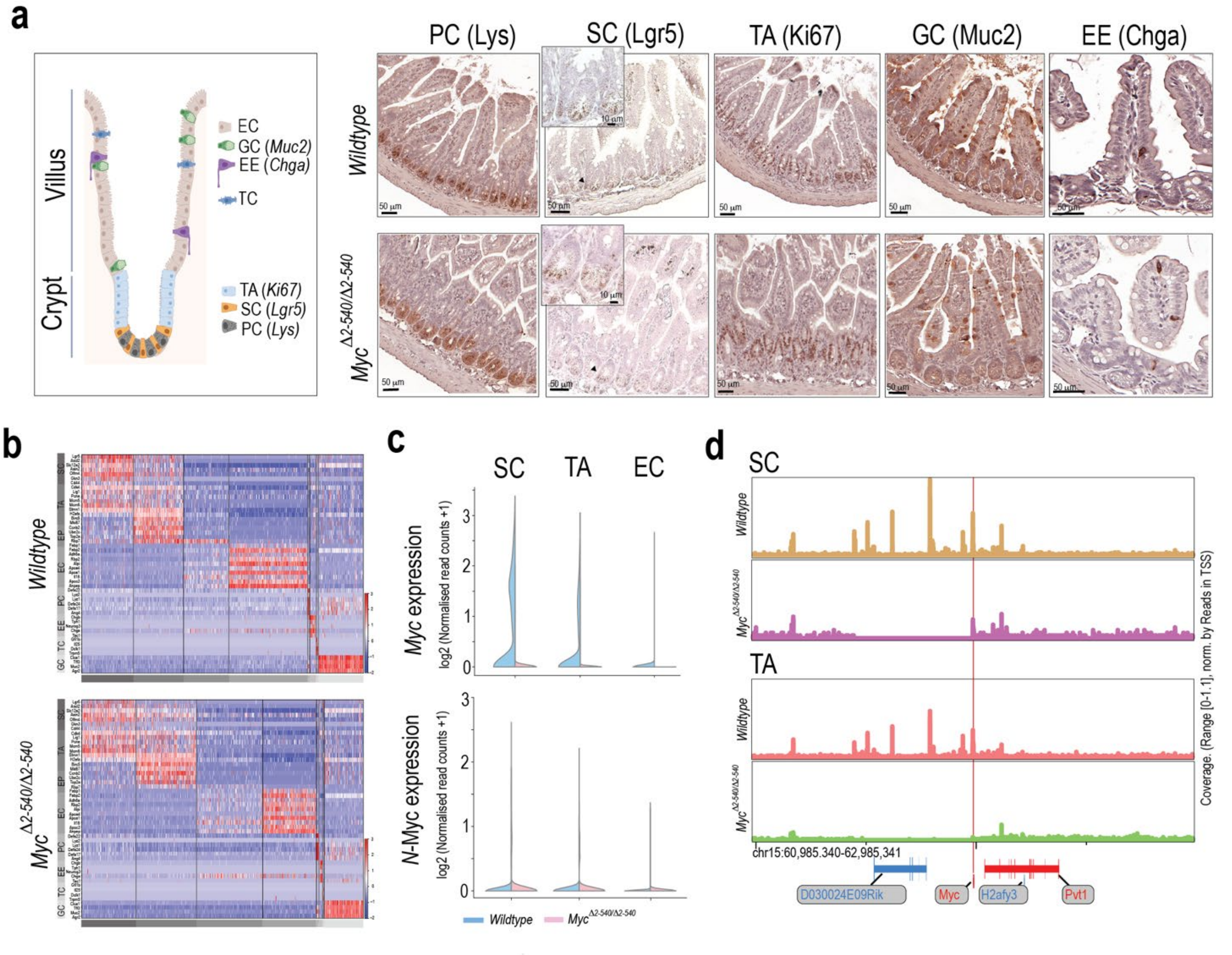


**Extended data Fig. 3**: ***$Myc^{\Delta 2\text{-}540/\Delta 2\text{-}540}$* intestine has a normal differentiation program during homeostasis a,** Similar pattern of cell-type specific marker expression in the wildtype and *$Myc^{\Delta 2\text{-}540/\Delta 2\text{-}540}$* intestine. IHC staining for Lysozyme (Lys), Ki67, Mucin2 (Muc2), Chromogranin A (Chga) and ISH staining for Lgr5 are shown. Left most panel: schematic of different cell types present within the intestinal epithelium. **b,** Heat map showing that all the major cell types of the intestine can be identified from single-cell RNA-seq analysis of intestinal crypts in both the wildtype and *$Myc^{\Delta 2\text{-}540/\Delta 2\text{-}540}$* mice. Clustering was done using top 2000 highly variable genes and cell type specific markers were according to Yum *et al* [46]. **c,** single-cell RNA-seq confirmed dramatic reduction in *Myc* expression within the SC and TA clusters. N-Myc expression was not detectable in both the wildtype and *$Myc^{\Delta 2\text{-}540/\Delta 2\text{-}540}$* mice. **d,** single-cell ATAC-seq analysis showing absence of compensatory changes in accessibility of *Myc* enhancer elements around the 538 kb enhancer region deleted in *$Myc^{\Delta 2\text{-}540/\Delta 2\text{-}540}$* mice.

PC: paneth cell, SC: stem cell; TA: transit amplifying cell, EE: enteroendocrine cell, TC: tuft cell; GC: goblet cell; EC: enterocyte; EP: enterocyte progenitor.

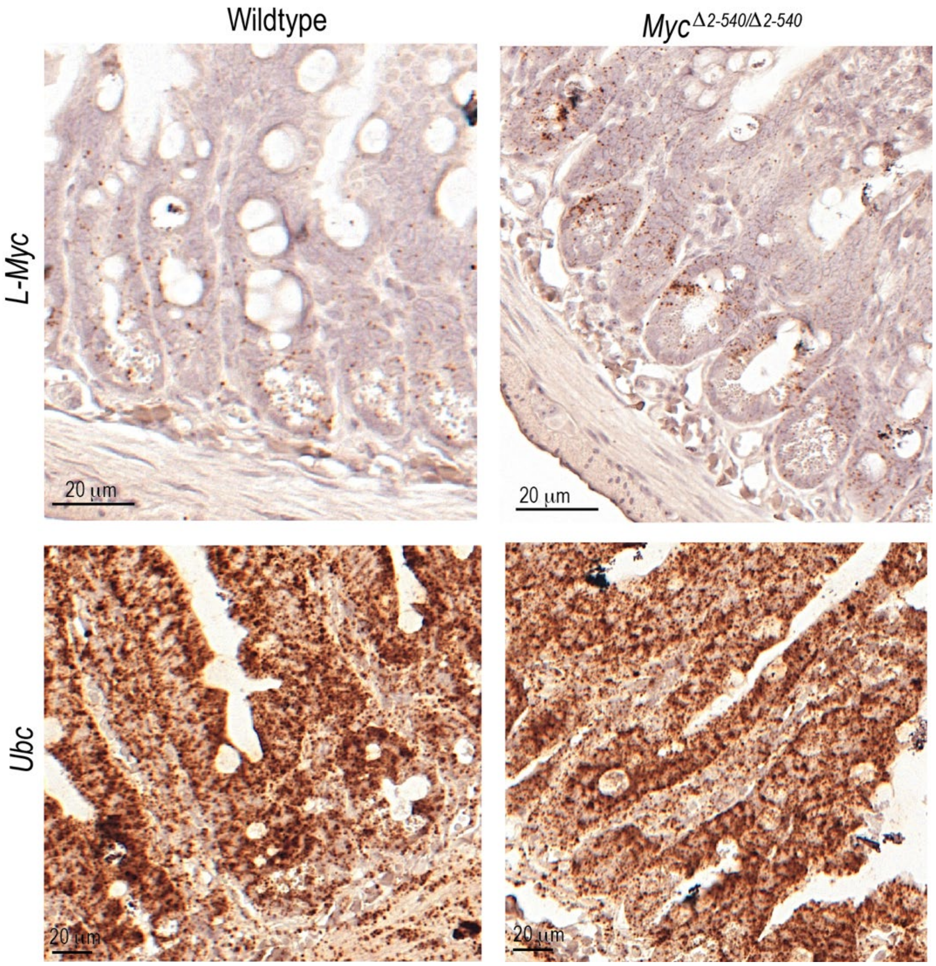


**Extended data Fig. 4**: ***Myc***$^{\Delta 2\text{-}540/\Delta 2\text{-}540}$ **mice have elevated *L-Myc* expression in the intestine during homeostasis**. ISH (RNAscope) showing increased expression of *L-Myc* transcripts in the intestine of unirradiated mice. Bright field microscope images are shown. *Ubc* transcripts were used as a positive control for RNA integrity of the sample.

#### Extended data Table 1

qPCR data for **Fig. 2c** showing loss of *Myc* induction after irradiation damage in intestine of $Myc^{\Delta 2\text{-}540/\Delta 2\text{-}540}$ mice.

| Genotype/ Treatment | Mouse ID | **Myc** | | **Actin** | | | | | |
|---|---|---|---|---|---|---|---|---|---|
| | | Cp | avg. Cp | Cp | avg. Cp | ΔCT | avg. ΔCT | (ΔΔCT) | Fold change |
| Wildtype (control) | 250/21 | 26,21 | 26,23 | 18,31 | 18,5 | 7,7 | 8,4 | 0,0 | 1,00 |
| | | 26,25 | | 18,52 | | | | | |
| | | 26,23 | | 18,65 | | | | | |
| Wildtype (control) | 255/21 | 26,05 | 26,25 | 18,04 | 18,0 | 8,2 | | | |
| | | 26,43 | | 18,07 | | | | | |
| | | 26,27 | | 18,03 | | | | | |
| Wildtype (control) | 221/21 | 27,92 | 27,84 | 18,68 | 18,7 | 9,2 | | | |
| | | 27,79 | | 18,64 | | | | | |
| | | 27,81 | | 18,69 | | | | | |
| Wildtype (3 dpi) | 211/21 | 26,8 | 26,80 | 19,93 | 20,0 | 6,8 | 7,1 | -1,3 | 2,46 |
| | | 26,72 | | 19,96 | | | | | |
| | | 26,87 | | 19,98 | | | | | |
| Wildtype (3 dpi) | 212/21 | 27,24 | 27,33 | 19,67 | 19,7 | 7,6 | | | |
| | | 27,27 | | 19,71 | | | | | |
| | | 27,49 | | 19,68 | | | | | |
| Wildtype (3 dpi) | 219/21 | 25,58 | 25,57 | 18,82 | 18,8 | 6,7 | | | |
| | | 25,52 | | 18,81 | | | | | |
| | | 25,61 | | 18,9 | | | | | |
| Wildtype (5 dpi) | 259/21 | 26,18 | 26,22 | 17,77 | 17,7 | 8,5 | 8,3 | -0,1 | 1,05 |
| | | 26,27 | | 17,75 | | | | | |
| | | 26,22 | | 17,71 | | | | | |
| Wildtype (5 dpi) | 260/21 | 25,74 | 25,77 | 17,53 | 17,5 | 8,2 | | | |
| | | 25,74 | | 17,49 | | | | | |
| | | 25,82 | | 17,55 | | | | | |
| Wildtype (5 dpi) | 263/21 | 26,06 | 26,05 | 17,84 | 17,9 | 8,2 | | | |
| | | 25,95 | | 17,85 | | | | | |
| | | 26,15 | | 17,89 | | | | | |
| $Myc^{D2\text{-}540/D2\text{-}540}$ (control) | 249/21 | 27,79 | 27,77 | 17,42 | 17,4 | 10,4 | 10,8 | 2,5 | 0,2 |
| | | 27,75 | | 17,23 | | | | | |
| | | 27,77 | | 17,42 | | | | | |
| $Myc^{D2\text{-}540/D2\text{-}540}$ (control) | 254/21 | 28,55 | 28,56 | 17,29 | 17,3 | 11,2 | | | |
| | | 28,55 | | 17,35 | | | | | |
| | | 28,59 | | 17,34 | | | | | |
| $Myc^{D2\text{-}540/D2\text{-}540}$ (3 dpi) | 210/21 | 29,63 | 29,59 | 18,79 | 19,2 | 10,4 | 10,9 | 2,6 | 0,17 |
| | | 29,55 | | 19,46 | | | | | |
| | | 29,59 | | 19,44 | | | | | |
| $Myc^{D2\text{-}540/D2\text{-}540}$ (3 dpi) | 214/21 | 31,07 | 31,05 | 19,52 | 19,5 | 11,5 | | | |
| | | 31,47 | | 19,51 | | | | | |
| | | 30,6 | | 19,53 | | | | | |
| $Myc^{D2\text{-}540/D2\text{-}540}$ (5 dpi) | 257/21 | 27,55 | 27,57 | 17,47 | 17,5 | 10,1 | 10,3 | 1,9 | 0,26 |
| | | 27,59 | | 17,45 | | | | | |
| | | 27,58 | | 17,54 | | | | | |
| $Myc^{D2\text{-}540/D2\text{-}540}$ (5 dpi) | 261/21 | 28,43 | 28,36 | 17,88 | 17,9 | 10,5 | | | |
| | | 28,18 | | 17,87 | | | | | |
| | | 28,48 | | 17,9 | | | | | |
| $Myc^{D2\text{-}540/D2\text{-}540}$ (5 dpi) | 262/21 | 28,26 | 28,20 | 17,9 | 17,9 | 10,3 | | | |
| | | 28,16 | | 17,86 | | | | | |
| | | 28,19 | | 17,96 | | | | | |

### Extended data Table 2

Mean expression (normalised read counts) of genes within 5Mb region around *Myc* whose expression can be detected in RNAseq data.

| Genotype | WT Irradiation (2dpi) | | | | | | | |
|---|---|---|---|---|---|---|---|---|
| Gene \ Cluster | SC | TA | EP | EC | PN | EE | TC | GC |
| **Myc** | 1,3751 | 1,1427 | 1,0695 | 0,0278 | 0,2051 | 0,2517 | 0,0658 | 0,3405 |
| **Fam49b** | 0,8123 | 0,9554 | 0,6943 | 0,8274 | 0,7664 | 1,3253 | 1,1568 | 0,7587 |
| **Nsmce2** | 0,7405 | 0,8266 | 0,8076 | 0,5472 | 0,4602 | 0,3452 | 0,1929 | 0,4250 |
| **Gsdmc4** | 0,0000 | 0,0007 | 0,1924 | 0,0030 | 0,0000 | 0,0000 | 0,0000 | 0,0000 |
| **Trib1** | 0,5683 | 0,4182 | 0,7972 | 2,8488 | 0,7276 | 0,9586 | 0,1743 | 0,7015 |
| **Lratd2** | 0,3587 | 0,2893 | 0,3828 | 0,3911 | 0,6420 | 0,2738 | 0,0593 | 0,4132 |
| **Gsdmc2** | 0,0043 | 0,0000 | 0,0326 | 0,0000 | 0,0000 | 0,0000 | 0,0000 | 0,0000 |
| **Gsdmc3** | 0,0000 | 0,0000 | 0,0000 | 0,0015 | 0,0000 | 0,0000 | 0,0000 | 0,0000 |
| **9930014A18Rik** | 0,0944 | 0,1419 | 0,0656 | 0,0313 | 0,0916 | 0,0103 | 0,0292 | 0,0621 |
| **Asap1** | 0,1937 | 0,1088 | 0,1695 | 2,7451 | 0,0492 | 0,4646 | 0,2107 | 0,2502 |
| **Pvt1** | 0,3914 | 0,3570 | 0,4085 | 0,0036 | 0,1863 | 0,1332 | 0,4529 | 0,1836 |
| **Gm38563** | 0,0580 | 0,0217 | 0,0819 | 0,0123 | 0,0000 | 0,0096 | 0,0000 | 0,0108 |
| **Gm36677** | 0,0000 | 0,0000 | 0,0020 | 0,0000 | 0,0056 | 0,0000 | 0,0000 | 0,0033 |

| Genotype | Myc $^{\Delta 2\text{-}540/\Delta 2\text{-}540}$ Irradiation (2dpi) | | | | | | | |
|---|---|---|---|---|---|---|---|---|
| Gene \ Cluster | SC | TA | EP | EC | PN | EE | TC | GC |
| **Myc** | 0,0073 | 0,0008 | 0,0006 | 0,0000 | 0,0946 | 0,0119 | 0,0110 | 0,0338 |
| **Fam49b** | 0,7468 | 0,8311 | 0,6830 | 0,7134 | 0,7450 | 1,9104 | 1,2513 | 0,5874 |
| **Nsmce2** | 0,7271 | 0,9548 | 0,6109 | 0,4270 | 0,4036 | 0,3725 | 0,3986 | 0,3318 |
| **Gsdmc4** | 0,0068 | 0,0044 | 0,0051 | 0,0000 | 0,0000 | 0,0000 | 0,0000 | 0,0000 |
| **Trib1** | 0,6981 | 0,6061 | 0,8725 | 0,6912 | 1,1123 | 1,0310 | 0,3294 | 0,8469 |
| **Lratd2** | 0,3487 | 0,2796 | 0,3569 | 0,4799 | 0,5875 | 0,3350 | 0,2758 | 0,4865 |
| **Gsdmc2** | 0,0021 | 0,0000 | 0,0000 | 0,0000 | 0,0000 | 0,0000 | 0,0000 | 0,0000 |
| **Gsdmc3** | 0,0000 | 0,0000 | 0,0000 | 0,0000 | 0,0000 | 0,0000 | 0,0000 | 0,0000 |
| **9930014A18Rik** | 0,0491 | 0,0612 | 0,0555 | 0,0326 | 0,0742 | 0,1084 | 0,0000 | 0,0529 |
| **Asap1** | 0,0910 | 0,0583 | 0,0903 | 0,4608 | 0,0525 | 0,5568 | 0,1970 | 0,1837 |
| **Pvt1** | 0,0321 | 0,0330 | 0,0282 | 0,0000 | 0,1941 | 0,0207 | 0,3674 | 0,0681 |
| **Gm38563** | 0,0088 | 0,0106 | 0,0231 | 0,0141 | 0,0158 | 0,0131 | 0,0000 | 0,0011 |
| **Gm36677** | 0,0018 | 0,0000 | 0,0017 | 0,0000 | 0,0000 | 0,0000 | 0,0000 | 0,0000 |

| Genotype | WT | | | | | | | |
|---|---|---|---|---|---|---|---|---|
| Gene \ Cluster | SC | TA | EP | EC | PN | EE | TC | GC |
| **Myc** | 0,9830 | 0,4696 | 0,1014 | 0,0214 | 0,0890 | 0,2675 | 0,1015 | 0,2698 |
| **Fam49b** | 0,8060 | 0,9269 | 0,6917 | 0,6948 | 0,6019 | 1,2693 | 1,4588 | 0,8092 |
| **Nsmce2** | 0,7467 | 0,9375 | 0,6353 | 0,5186 | 0,3544 | 0,4969 | 0,4860 | 0,5632 |
| **Gsdmc4** | 0,7260 | 2,5320 | 8,7972 | 0,6179 | 0,3366 | 0,3846 | 0,4823 | 0,5642 |
| **Trib1** | 0,4437 | 0,2591 | 0,2298 | 2,4042 | 0,8430 | 0,7727 | 0,2732 | 0,5534 |
| **Lratd2** | 0,2913 | 0,2322 | 0,3065 | 0,5813 | 0,7535 | 0,2815 | 0,1378 | 0,4596 |
| **Gsdmc2** | 0,2066 | 0,8604 | 2,1084 | 0,1485 | 0,0441 | 0,1413 | 0,1002 | 0,1857 |
| **Gsdmc3** | 0,1256 | 0,4439 | 1,1678 | 0,0926 | 0,0093 | 0,0425 | 0,0768 | 0,0986 |
| **9930014A18Rik** | 0,1183 | 0,0982 | 0,0702 | 0,0192 | 0,0854 | 0,0781 | 0,0990 | 0,0534 |
| **Asap1** | 0,0967 | 0,1043 | 0,0783 | 1,0038 | 0,0475 | 0,3087 | 0,0894 | 0,0583 |
| **Pvt1** | 0,0103 | 0,0081 | 0,0000 | 0,0000 | 0,0435 | 0,0405 | 0,1083 | 0,0508 |
| **Gm38563** | 0,0069 | 0,0022 | 0,0188 | 0,0030 | 0,0000 | 0,0059 | 0,0058 | 0,0030 |
| **Gm36677** | 0,0000 | 0,0006 | 0,0000 | 0,0000 | 0,0000 | 0,0035 | 0,0000 | 0,0016 |

| Genotype | $Myc^{\Delta 2\text{-}540/\Delta 2\text{-}540}$ | | | | | | | |
|---|---|---|---|---|---|---|---|---|
| Gene \ Cluster | SC | TA | EP | EC | PN | EE | TC | GC |
| **Myc** | 0,0276 | 0,0067 | 0,0029 | 0,0000 | 0,0936 | 0,0356 | 0,0978 | 0,1064 |
| **Fam49b** | 0,8179 | 0,9545 | 0,8399 | 0,8322 | 1,3815 | 1,4700 | 1,4192 | 0,8691 |
| **Nsmce2** | 0,7101 | 0,8481 | 0,6575 | 0,4536 | 0,6049 | 0,3030 | 0,7063 | 0,5248 |
| **Gsdmc4** | 0,2074 | 0,4029 | 1,7180 | 0,0893 | 0,2387 | 0,0304 | 0,0185 | 0,1265 |
| **Trib1** | 0,5504 | 0,3975 | 0,4556 | 1,8144 | 1,0253 | 0,6644 | 0,2765 | 0,6771 |
| **Lratd2** | 0,3485 | 0,2994 | 0,4317 | 0,5305 | 1,1676 | 0,4156 | 0,2063 | 0,6753 |
| **Gsdmc2** | 0,0475 | 0,1277 | 0,4204 | 0,0209 | 0,0000 | 0,0095 | 0,0000 | 0,0449 |
| **Gsdmc3** | 0,0192 | 0,0461 | 0,1719 | 0,0129 | 0,0000 | 0,0063 | 0,0000 | 0,0215 |
| **9930014A18Rik** | 0,0841 | 0,0753 | 0,0541 | 0,0361 | 0,0693 | 0,0256 | 0,0372 | 0,0546 |
| **Asap1** | 0,0599 | 0,0673 | 0,0954 | 0,7649 | 0,1170 | 0,4795 | 0,1344 | 0,0699 |
| **Pvt1** | 0,0007 | 0,0000 | 0,0000 | 0,0000 | 0,2682 | 0,0000 | 0,1005 | 0,0354 |
| **Gm38563** | 0,0107 | 0,0028 | 0,0252 | 0,0146 | 0,0000 | 0,0000 | 0,0000 | 0,0009 |
| **Gm36677** | 0,0018 | 0,0002 | 0,0018 | 0,0000 | 0,0000 | 0,0069 | 0,0000 | 0,0070 |

**SC:** Stem cell; **TA:** Transit amplifying cell; **EP:** Enterocyte progenitor; **EC:** Enterocyte;
**PN:** Paneth cells; **EE:** Enteroendocrine; **TC:** Tuft cell; **GC:** Goblet cell

### Extended data Table 3

GSEA results for **a**: Gene Ontology Biological Process (GOBP) terms and **b**: Molecular Signatures Database (MSigDB) Hallmark gene set within the stem cell (SC) population

**a**: Top 100 GSEA results for Gene Ontology Biological Process (GOBP) terms in SC

| ID | setSize | ES | NES | pvalue | p.adjust |
|---|---|---|---|---|---|
| GOBP_RIBOSOME_BIOGENESIS | 290 | 0,658550092 | 1,748875109 | 3,65358E-08 | 0,000173326 |
| GOBP_NCRNA_PROCESSING | 356 | 0,612548346 | 1,655358234 | 7,98566E-08 | 0,00018942 |
| GOBP_RRNA_METABOLIC_PROCESS | 238 | 0,66298767 | 1,752693838 | 8,26613E-07 | 0,001307151 |
| GOBP_RIBONUCLEOPROTEIN_COMPLEX_BIOGENESIS | 402 | 0,569923507 | 1,547174243 | 1,70293E-06 | 0,002019677 |
| GOBP_FOREBRAIN_REGIONALIZATION | 11 | -0,962809924 | -1,462865026 | 2,61593E-05 | 0,024819955 |
| GOBP_RESPONSE_TO_ETHANOL | 48 | -0,858299916 | -1,583065747 | 8,63348E-05 | 0,068262039 |
| GOBP_REGULATION_OF_CAMP_DEPENDENT_PROTEIN_KINASE_ ACTIVITY | 12 | 0,944239095 | 1,709317435 | 0,000152429 | 0,103303574 |
| GOBP_COLLATERAL_SPROUTING | 24 | 0,882236309 | 1,718170937 | 0,000537363 | 0,266984071 |
| GOBP_REGULATION_OF_STORE_OPERATED_CALCIUM_ENTRY | 15 | 0,925061895 | 1,667304223 | 0,000459753 | 0,266984071 |
| GOBP_MITOCHONDRIAL_ELECTRON_TRANSPORT_CYTOCHROME_C_ TO_OXYGEN | 12 | -0,933624623 | -1,42798416 | 0,000562783 | 0,266984071 |
| GOBP_TISSUE_REGENERATION | 40 | 0,828546309 | 1,719127625 | 0,000730605 | 0,315089959 |
| GOBP_ORGANELLE_FUSION | 125 | -0,737686936 | -1,542249147 | 0,000856667 | 0,330401182 |
| GOBP_SYNAPTIC_TRANSMISSION_GABAERGIC | 40 | -0,848941741 | -1,516608662 | 0,0009054 | 0,330401182 |
| GOBP_EXOCYTIC_PROCESS | 79 | -0,785367193 | -1,555331387 | 0,001070024 | 0,342574808 |
| GOBP_CELL_PROLIFERATION_IN_HINDBRAIN | 11 | -0,93810454 | -1,425328395 | 0,001083183 | 0,342574808 |
| GOBP_REGULATION_OF_CARDIOCYTE_DIFFERENTIATION | 23 | 0,877266861 | 1,698368028 | 0,001212015 | 0,359362563 |
| GOBP_ORGANELLE_MEMBRANE_FUSION | 96 | -0,752327072 | -1,537413901 | 0,001396324 | 0,368008948 |
| GOBP_HISTAMINE_TRANSPORT | 13 | -0,923783555 | -1,404543394 | 0,001327842 | 0,368008948 |
| GOBP_EPIBOLY | 26 | 0,852708603 | 1,673084696 | 0,002316516 | 0,453904369 |
| GOBP_REGULATION_OF_ENDOCRINE_PROCESS | 30 | 0,83783291 | 1,67224599 | 0,002871371 | 0,453904369 |
| GOBP_STORE_OPERATED_CALCIUM_ENTRY | 21 | 0,873004873 | 1,669457625 | 0,003228197 | 0,453904369 |
| GOBP_REGULATION_OF_COLLATERAL_SPROUTING | 17 | 0,895109491 | 1,637747028 | 0,002816398 | 0,453904369 |
| GOBP_RESPONSE_TO_ALKALOID | 62 | -0,794438577 | -1,532266865 | 0,003108423 | 0,453904369 |
| GOBP_SYNAPTIC_TRANSMISSION_GLUTAMATERGIC | 68 | -0,782406351 | -1,530198127 | 0,003313145 | 0,453904369 |
| GOBP_MEMBRANE_FUSION | 121 | -0,712930597 | -1,488932048 | 0,003381477 | 0,453904369 |
| GOBP_REGULATION_OF_SYNAPTIC_TRANSMISSION_GABAERGIC | 27 | -0,864930766 | -1,465824858 | 0,003444468 | 0,453904369 |
| GOBP_ENDOCARDIAL_CUSHION_DEVELOPMENT | 32 | -0,846967376 | -1,464562106 | 0,00327465 | 0,453904369 |
| GOBP_TRNA_METABOLIC_PROCESS | 168 | 0,570217967 | 1,433843375 | 0,003046715 | 0,453904369 |
| GOBP_REGULATION_OF_FEEDING_BEHAVIOR | 19 | -0,89028592 | -1,424777749 | 0,003380959 | 0,453904369 |
| GOBP_REGULATION_OF_CHROMOSOME_ORGANIZATION | 199 | 0,550489057 | 1,424623387 | 0,00232636 | 0,453904369 |
| GOBP_DNA_CONFORMATION_CHANGE | 217 | 0,544034997 | 1,411288401 | 0,0023778 | 0,453904369 |
| GOBP_REGULATION_OF_SKELETAL_MUSCLE_CELL_ DIFFERENTIATION | 15 | -0,905242444 | -1,411053529 | 0,002932228 | 0,453904369 |
| GOBP_PONS_DEVELOPMENT | 11 | -0,926979102 | -1,408424731 | 0,003187205 | 0,453904369 |
| GOBP_POSITIVE_REGULATION_OF_FIBROBLAST_MIGRATION | 14 | -0,9103963 | -1,400670205 | 0,002535316 | 0,453904369 |
| GOBP_REGULATION_OF_CELL_PROJECTION_ASSEMBLY | 174 | 0,551543756 | 1,396472694 | 0,003120463 | 0,453904369 |
| GOBP_NCRNA_METABOLIC_PROCESS | 467 | 0,462521114 | 1,250790187 | 0,003132959 | 0,453904369 |
| GOBP_POSITIVE_REGULATION_OF_SYNAPTIC_TRANSMISSION_ GLUTAMATERGIC | 18 | -0,893638323 | -1,408371698 | 0,003665294 | 0,469950176 |
| GOBP_NEURAL_NUCLEUS_DEVELOPMENT | 17 | -0,894824788 | -1,400057502 | 0,00405473 | 0,506200976 |
| GOBP_RIBOSOMAL_LARGE_SUBUNIT_BIOGENESIS | 68 | 0,707837905 | 1,570881809 | 0,004164034 | 0,506517338 |
| GOBP_ORGANIC_CATION_TRANSPORT | 27 | -0,860478062 | -1,458278723 | 0,004739797 | 0,562139873 |
| GOBP_NEGATIVE_REGULATION_OF_PROTEIN_SERINE_THREONINE_ KINASE_ACTIVITY | 96 | 0,632614146 | 1,480140557 | 0,005167477 | 0,597914935 |
| GOBP_BERGMANN_GLIAL_CELL_DIFFERENTIATION | 10 | -0,925352495 | -1,402334992 | 0,005295894 | 0,598183783 |
| GOBP_STRESS_FIBER_ASSEMBLY | 101 | 0,626232511 | 1,47982169 | 0,006261854 | 0,670037214 |
| GOBP_ACTIN_FILAMENT_BUNDLE_ORGANIZATION | 153 | 0,581044115 | 1,422486232 | 0,006306761 | 0,670037214 |
| GOBP_CATECHOLAMINE_UPTAKE | 12 | -0,907157996 | -1,387503304 | 0,006355749 | 0,670037214 |
| GOBP_PROTEIN_KINASE_A_SIGNALING | 24 | 0,837767571 | 1,631567279 | 0,007103319 | 0,673957536 |
| GOBP_DETECTION_OF_CHEMICAL_STIMULUS_INVOLVED_IN_ SENSORY_PERCEPTION_OF_TASTE | 10 | 0,912176344 | 1,631145783 | 0,007019977 | 0,673957536 |
| GOBP_ENDOCRINE_HORMONE_SECRETION | 37 | 0,789342028 | 1,607241011 | 0,006994431 | 0,673957536 |
| GOBP_NEGATIVE_REGULATION_OF_PROTEIN_KINASE_B_SIGNALING | 40 | 0,774484101 | 1,606955457 | 0,006963458 | 0,673957536 |
| GOBP_FATTY_ACID_CATABOLIC_PROCESS | 90 | -0,724722196 | -1,471730686 | 0,007246441 | 0,673957536 |
| GOBP_RESPONSE_TO_CAFFEINE | 11 | -0,918881032 | -1,396120762 | 0,007387393 | 0,673957536 |
| GOBP_VASCULAR_ENDOTHELIAL_CELL_PROLIFERATION | 18 | -0,885129878 | -1,394962411 | 0,007238673 | 0,673957536 |

| | | | | | |
|---|---|---|---|---|---|
| GOBP_REGULATION_OF_GLYCOPROTEIN_METABOLIC_PROCESS | 39 | 0,780751864 | 1,606507076 | 0,007869981 | 0,674272352 |
| GOBP_SECRETORY_GRANULE_LOCALIZATION | 16 | 0,874250407 | 1,594056905 | 0,008022831 | 0,674272352 |
| GOBP_FATTY_ACID_BETA_OXIDATION | 67 | -0,756510902 | -1,476655229 | 0,007953028 | 0,674272352 |
| GOBP_STEM_CELL_DIFFERENTIATION | 163 | 0,555806501 | 1,397508842 | 0,007795881 | 0,674272352 |
| GOBP_POSITIVE_REGULATION_OF_BEHAVIOR | 20 | -0,86451566 | -1,389908314 | 0,008101502 | 0,674272352 |
| GOBP_REGULATION_OF_ACTOMYOSIN_STRUCTURE_ORGANIZATION | 96 | 0,615803861 | 1,44080918 | 0,008775794 | 0,717799448 |
| GOBP_LUTEINIZATION | 10 | 0,905256665 | 1,618772073 | 0,009533356 | 0,72074303 |
| GOBP_NEGATIVE_REGULATION_OF_B_CELL_ACTIVATION | 24 | 0,830991608 | 1,618370969 | 0,009650681 | 0,72074303 |
| GOBP_STRESS_RESPONSE_TO_METAL_ION | 10 | 0,904819237 | 1,617989868 | 0,009533356 | 0,72074303 |
| GOBP_PHOSPHOLIPASE_C_ACTIVATING_G_PROTEIN_COUPLED_ RECEPTOR_SIGNALING_PATHWAY | 36 | 0,779344892 | 1,592159165 | 0,009839026 | 0,72074303 |
| GOBP_RIBOSOMAL_SMALL_SUBUNIT_BIOGENESIS | 71 | 0,671022882 | 1,510329458 | 0,010287555 | 0,72074303 |
| GOBP_NEGATIVE_REGULATION_OF_DEVELOPMENTAL_GROWTH | 101 | 0,612775827 | 1,448022809 | 0,010341029 | 0,72074303 |
| GOBP_MEMBRANE_DOCKING | 75 | -0,734363414 | -1,444040682 | 0,010197418 | 0,72074303 |
| GOBP_HISTAMINE_SECRETION | 11 | -0,916212303 | -1,392065975 | 0,010482983 | 0,72074303 |
| GOBP_RESPONSE_TO_ISOQUINOLINE_ALKALOID | 15 | -0,886310637 | -1,381543432 | 0,009903249 | 0,72074303 |
| GOBP_SPONTANEOUS_SYNAPTIC_TRANSMISSION | 12 | -0,902180997 | -1,379890956 | 0,009459903 | 0,72074303 |
| GOBP_HEAT_GENERATION | 14 | -0,892196717 | -1,372669638 | 0,009304537 | 0,72074303 |
| GOBP_NEGATIVE_REGULATION_OF_RESPONSE_TO_FOOD | 11 | 0,900212058 | 1,62943945 | 0,011409361 | 0,721680094 |
| GOBP_POSITIVE_REGULATION_OF_DOUBLE_STRAND_ BREAK_REPAIR | 64 | 0,686292636 | 1,508312371 | 0,011122968 | 0,721680094 |
| GOBP_DEFENSE_RESPONSE_TO_GRAM_POSITIVE_BACTERIUM | 82 | 0,621401246 | 1,450676553 | 0,011015937 | 0,721680094 |
| GOBP_DNA_PACKAGING | 163 | 0,545480345 | 1,371544962 | 0,011285973 | 0,721680094 |
| GOBP_RNA_MODIFICATION | 139 | 0,56707857 | 1,364611726 | 0,010873809 | 0,721680094 |
| GOBP_RESPONSE_TO_WOUNDING | 374 | 0,448083608 | 1,215473943 | 0,010964451 | 0,721680094 |
| GOBP_MATURATION_OF_SSU_RRNA | 49 | 0,699682355 | 1,503094978 | 0,012258856 | 0,745589932 |
| GOBP_NUCLEUS_ORGANIZATION | 123 | 0,596525633 | 1,426123978 | 0,011987758 | 0,745589932 |
| GOBP_NEGATIVE_REGULATION_OF_GROWTH | 211 | 0,504049271 | 1,303500263 | 0,012174697 | 0,745589932 |
| GOBP_ORGAN_OR_TISSUE_SPECIFIC_IMMUNE_RESPONSE | 36 | 0,765991301 | 1,564878505 | 0,013082146 | 0,760171849 |
| GOBP_GLUTAMATE_RECEPTOR_SIGNALING_PATHWAY | 30 | -0,829019492 | -1,425856513 | 0,013117331 | 0,760171849 |
| GOBP_MONOCARBOXYLIC_ACID_CATABOLIC_PROCESS | 105 | -0,69306393 | -1,424831096 | 0,012996818 | 0,760171849 |
| GOBP_PULMONARY_VALVE_DEVELOPMENT | 15 | -0,881865608 | -1,374614709 | 0,013139564 | 0,760171849 |
| GOBP_REGULATION_OF_BEHAVIOR | 53 | -0,770671102 | -1,452650973 | 0,013310726 | 0,760796174 |
| GOBP_GANGLION_DEVELOPMENT | 10 | 0,897325122 | 1,604588958 | 0,013651692 | 0,770995543 |
| GOBP_POSITIVE_REGULATION_OF_AXONOGENESIS | 82 | 0,610461444 | 1,425137316 | 0,013990863 | 0,780080481 |
| GOBP_VOCALIZATION_BEHAVIOR | 11 | -0,911417437 | -1,384780797 | 0,014141425 | 0,780080481 |
| GOBP_GASTRULATION_WITH_MOUTH_FORMING_SECOND | 31 | -0,824920299 | -1,426083326 | 0,014890451 | 0,811957482 |
| GOBP_RESPONSE_TO_LEPTIN | 17 | 0,861702792 | 1,57662409 | 0,01525763 | 0,822524956 |
| GOBP_POSITIVE_REGULATION_OF_CARDIOCYTE_DIFFERENTIATION | 14 | 0,869223062 | 1,571543887 | 0,015437963 | 0,822895454 |
| GOBP_NEGATIVE_REGULATION_OF_B_CELL_PROLIFERATION | 14 | 0,868441078 | 1,57013007 | 0,016105544 | 0,840677529 |
| GOBP_POSITIVE_REGULATION_OF_EXOCYTOSIS | 76 | -0,71299882 | -1,404690385 | 0,016125981 | 0,840677529 |
| GOBP_SEROTONIN_TRANSPORT | 19 | -0,867123372 | -1,387709339 | 0,016414268 | 0,846405284 |
| GOBP_SPHINGOLIPID_MEDIATED_SIGNALING_PATHWAY | 10 | 0,888572643 | 1,588937851 | 0,017321465 | 0,855969061 |
| GOBP_POSITIVE_REGULATION_OF_MUSCLE_HYPERTROPHY | 31 | 0,788484075 | 1,572350077 | 0,016870821 | 0,855969061 |
| GOBP_POSITIVE_REGULATION_OF_GLUTAMATE_SECRETION | 10 | -0,909324957 | -1,378045894 | 0,017318461 | 0,855969061 |
| GOBP_MAINTENANCE_OF_CELL_NUMBER | 141 | 0,534396588 | 1,281576452 | 0,017218113 | 0,855969061 |
| GOBP_RESPONSE_TO_INTERFERON_BETA | 36 | 0,749806304 | 1,531813437 | 0,017949837 | 0,870609143 |
| GOBP_DEVELOPMENTAL_CELL_GROWTH | 213 | 0,494955022 | 1,281882786 | 0,017984759 | 0,870609143 |
| GOBP_VESICLE_FUSION_TO_PLASMA_MEMBRANE | 21 | -0,850129801 | -1,370634523 | 0,018486694 | 0,885867416 |
| GOBP_NEGATIVE_REGULATION_OF_LIPID_BIOSYNTHETIC_PROCESS | 48 | 0,691541771 | 1,488526428 | 0,019143752 | 0,901543161 |

**b:** GSEA results for Molecular Signatures Database (MSigDB) Hallmark gene set within the stem cell (SC) population

| ID | setSize | ES | NES | pvalue | p.adjust |
|---|---|---|---|---|---|
| HALLMARK_MYC_TARGETS_V1 | 191 | 0,629989994 | 1,617036414 | 6,51932E-05 | 0,00325966 |
| HALLMARK_E2F_TARGETS | 196 | 0,604678817 | 1,549167059 | 0,000204117 | 0,005102928 |
| HALLMARK_G2M_CHECKPOINT | 187 | 0,56019513 | 1,427728245 | 0,002329621 | 0,038827014 |
| HALLMARK_UNFOLDED_PROTEIN_RESPONSE | 108 | 0,544339096 | 1,336227721 | 0,045256467 | 0,565705839 |
| HALLMARK_NOTCH_SIGNALING | 30 | -0,767612842 | -1,326665981 | 0,0672 | 0,567589046 |
| HALLMARK_ALLOGRAFT_REJECTION | 118 | 0,505265771 | 1,2403318 | 0,068110685 | 0,567589046 |
| HALLMARK_MYC_TARGETS_V2 | 58 | 0,546791727 | 1,227240337 | 0,122743682 | 0,853156477 |
| HALLMARK_IL6_JAK_STAT3_SIGNALING | 66 | 0,521058923 | 1,201631608 | 0,140684411 | 0,853156477 |
| HALLMARK_MTORC1_SIGNALING | 190 | 0,438170419 | 1,119672525 | 0,153568166 | 0,853156477 |
| HALLMARK_GLYCOLYSIS | 181 | 0,424290185 | 1,079557752 | 0,214765101 | 1 |
| HALLMARK_APICAL_JUNCTION | 155 | -0,498927655 | -1,053244648 | 0,364497041 | 1 |
| HALLMARK_HEDGEHOG_SIGNALING | 27 | 0,521308146 | 1,045239129 | 0,44386423 | 1 |
| HALLMARK_ESTROGEN_RESPONSE_LATE | 170 | -0,488337853 | -1,036109921 | 0,387706856 | 1 |
| HALLMARK_EPITHELIAL_MESENCHYMAL_TRANSITION | 125 | -0,495397238 | -1,025828024 | 0,426650367 | 1 |
| HALLMARK_P53_PATHWAY | 184 | 0,391244357 | 0,994992902 | 0,492957746 | 1 |
| HALLMARK_TNFA_SIGNALING_VIA_NFKB | 176 | -0,467386547 | -0,993564976 | 0,474616293 | 1 |
| HALLMARK_HEME_METABOLISM | 164 | -0,460323243 | -0,976439053 | 0,516470588 | 1 |
| HALLMARK_INTERFERON_ALPHA_RESPONSE | 88 | 0,401891113 | 0,957887826 | 0,562211982 | 1 |
| HALLMARK_INFLAMMATORY_RESPONSE | 134 | -0,449637625 | -0,938388548 | 0,594011976 | 1 |
| HALLMARK_KRAS_SIGNALING_UP | 149 | -0,444930137 | -0,935780017 | 0,611442193 | 1 |
| HALLMARK_HYPOXIA | 167 | 0,368521813 | 0,931192661 | 0,696774194 | 1 |
| HALLMARK_REACTIVE_OXIGEN_SPECIES_PATHWAY | 47 | -0,501215596 | -0,930949568 | 0,592261905 | 1 |
| HALLMARK_SPERMATOGENESIS | 85 | -0,464968504 | -0,928324713 | 0,597938144 | 1 |
| HALLMARK_KRAS_SIGNALING_DN | 103 | -0,455260854 | -0,926086788 | 0,602996255 | 1 |
| HALLMARK_PEROXISOME | 93 | 0,371268063 | 0,891198777 | 0,719457014 | 1 |
| HALLMARK_XENOBIOTIC_METABOLISM | 164 | -0,415116923 | -0,88054727 | 0,754117647 | 1 |
| HALLMARK_COMPLEMENT | 148 | 0,352084364 | 0,874274345 | 0,853658537 | 1 |
| HALLMARK_MYOGENESIS | 138 | -0,407429887 | -0,853362887 | 0,776849642 | 1 |
| HALLMARK_DNA_REPAIR | 146 | 0,341920718 | 0,847806069 | 0,909090909 | 1 |
| HALLMARK_BILE_ACID_METABOLISM | 91 | -0,411048396 | -0,823830266 | 0,794155019 | 1 |
| HALLMARK_WNT_BETA_CATENIN_SIGNALING | 38 | -0,458565426 | -0,821563352 | 0,755007704 | 1 |
| HALLMARK_APICAL_SURFACE | 31 | -0,472962993 | -0,820781512 | 0,773734177 | 1 |
| HALLMARK_PANCREAS_BETA_CELLS | 34 | -0,462453055 | -0,817861859 | 0,788018433 | 1 |
| HALLMARK_OXIDATIVE_PHOSPHORYLATION | 194 | -0,364405404 | -0,782237564 | 0,923344948 | 1 |
| HALLMARK_INTERFERON_GAMMA_RESPONSE | 158 | -0,367948223 | -0,777956912 | 0,90887574 | 1 |
| HALLMARK_ESTROGEN_RESPONSE_EARLY | 180 | 0,304025836 | 0,77549692 | 0,993421053 | 1 |
| HALLMARK_UV_RESPONSE_UP | 138 | -0,36819633 | -0,771188106 | 0,918854415 | 1 |
| HALLMARK_ANGIOGENESIS | 25 | -0,449082603 | -0,756062647 | 0,874587459 | 1 |
| HALLMARK_UV_RESPONSE_DN | 129 | -0,361392341 | -0,751194513 | 0,943030303 | 1 |
| HALLMARK_CHOLESTEROL_HOMEOSTASIS | 69 | 0,318934798 | 0,74663944 | 0,962406015 | 1 |
| HALLMARK_PROTEIN_SECRETION | 92 | -0,372577522 | -0,74615195 | 0,921219822 | 1 |
| HALLMARK_IL2_STAT5_SIGNALING | 163 | -0,3501763 | -0,742019279 | 0,958726415 | 1 |
| HALLMARK_APOPTOSIS | 141 | 0,294103961 | 0,729657744 | 0,993710692 | 1 |
| HALLMARK_COAGULATION | 100 | -0,352127547 | -0,712768985 | 0,964512041 | 1 |
| HALLMARK_ADIPOGENESIS | 186 | -0,332032153 | -0,711462588 | 0,987209302 | 1 |
| HALLMARK_FATTY_ACID_METABOLISM | 144 | -0,326410439 | -0,683967461 | 0,989247312 | 1 |
| HALLMARK_ANDROGEN_RESPONSE | 90 | -0,336874246 | -0,676126289 | 0,983418367 | 1 |
| HALLMARK_PI3K_AKT_MTOR_SIGNALING | 95 | 0,266703183 | 0,641066998 | 1 | 1 |
| HALLMARK_MITOTIC_SPINDLE | 195 | 0,237309172 | 0,608580481 | 1 | 1 |
| HALLMARK_TGF_BETA_SIGNALING | 50 | -0,323769188 | -0,608487037 | 0,988522238 | 1 |

### Extended data Table 4

Mean expression (Normalized read counts) of MYC targets (**a**) and MYC family (**b**) in Stem (SC), Transit amplifying (TA) and Enterocyte (EC) cell clusters.

**a**: Mean expression (Normalised read counts) of MYC targets

Non-irradiated

| **Genotype** | **Myc$^{\Delta 2\text{-}540/\Delta 2\text{-}540}$** | | | **WT** | | | **Fold change (log2) Myc$^{\Delta 2\text{-}540/\Delta 2\text{-}540}$ *vs* WT** | | |
|---|---|---|---|---|---|---|---|---|---|
| **Gene / Cluster** | SC | TA | EC | SC | TA | EC | SC | TA | EC |
| **Atic** | 1,323163 | 1,182003 | 0,133417 | 1,990572 | 1,663091 | 0,096464 | -0,421825 | -0,363395 | 0,622574 |
| **Ctps** | 0,380310 | 0,310912 | 0,014385 | 0,437887 | 0,336974 | 0,008092 | 0,011619 | 0,060205 | 0,968191 |
| **Ctps2** | 0,204032 | 0,193485 | 0,075192 | 0,277964 | 0,235979 | 0,085141 | -0,203139 | -0,085990 | 0,077607 |
| **Nme1** | 6,482093 | 6,183938 | 1,664926 | 7,618751 | 7,853556 | 1,061065 | -0,139171 | -0,292126 | 0,773079 |
| **Nme2** | 11,887809 | 11,303824 | 6,201468 | 21,152828 | 20,261341 | 5,251464 | -0,848683 | -0,875157 | 0,402900 |
| **Ppat** | 0,477735 | 0,227627 | 0,011818 | 0,703715 | 0,369139 | 0,006735 | -0,344374 | -0,550039 | 0,764891 |
| **Ahcy** | 0,495351 | 0,490284 | 0,137489 | 0,448922 | 0,459889 | 0,123550 | 0,381237 | 0,280091 | 0,170593 |
| **Shmt2** | 0,212686 | 0,209862 | 0,019161 | 0,442318 | 0,382560 | 0,027877 | -0,825563 | -0,704380 | -0,474464 |
| **Aldh7a1** | 0,180028 | 0,148260 | 0,114803 | 0,205796 | 0,161823 | 0,094130 | 0,026685 | 0,068882 | 0,043655 |
| **Glo1** | 2,708696 | 2,920935 | 2,683761 | 2,502534 | 3,003896 | 2,396569 | 0,306195 | 0,085357 | 0,203188 |
| **Pck1** | 0,014109 | 0,007014 | 0,212210 | 0,015463 | 0,006334 | 0,495310 | 0,245017 | 0,257243 | -1,619664 |
| **Srm** | 0,298472 | 0,228423 | 0,053431 | 0,623399 | 0,518650 | 0,021299 | -0,845636 | -1,021513 | 1,341007 |
| **Acp2** | 0,209667 | 0,172554 | 0,342485 | 0,189284 | 0,129375 | 0,368529 | 0,384740 | 0,572862 | -0,092795 |
| **Dph5** | 0,311762 | 0,235665 | 0,131833 | 0,449542 | 0,390557 | 0,103339 | -0,340826 | -0,608036 | 0,425426 |
| **Hexa** | 1,378220 | 1,552756 | 2,327163 | 1,418477 | 1,721942 | 1,989858 | 0,160619 | 0,042336 | 0,254349 |
| **Dhrs3** | 0,357493 | 0,338971 | 0,610152 | 0,289160 | 0,297051 | 0,844236 | 0,564422 | 0,393854 | -0,443727 |
| **Slc2a6** | 0,000000 | 0,000000 | 0,000000 | 0,000000 | 0,000000 | 0,000000 | 0,000000 | 0,000000 | 0,000000 |
| **Hspe1** | 12,263428 | 12,141154 | 2,096624 | 15,946391 | 15,795172 | 1,468087 | -0,351759 | -0,364067 | 0,596828 |
| **Prmt3** | 0,198536 | 0,141461 | 0,023046 | 0,330351 | 0,245897 | 0,014979 | -0,514066 | -0,616142 | 0,673959 |
| **Psme2** | 5,378239 | 5,791747 | 6,110400 | 6,407434 | 8,237217 | 6,091765 | -0,128494 | -0,449576 | 0,017315 |
| **Rab11fip1** | 0,822493 | 0,765249 | 1,576507 | 0,654023 | 0,586995 | 2,160854 | 0,577359 | 0,581989 | -0,448909 |
| **Uevld** | 0,085805 | 0,241918 | 0,134044 | 0,104423 | 0,225564 | 0,140348 | -0,036466 | 0,283438 | -0,112245 |
| **Usp2** | 0,249178 | 0,298111 | 0,791815 | 0,271799 | 0,421632 | 0,735360 | 0,027213 | -0,313221 | 0,119127 |
| **Tnfaip8l1** | 0,447765 | 0,583085 | 0,135526 | 0,446440 | 0,571022 | 0,067751 | 0,277596 | 0,231103 | 0,960060 |
| **Gulp1** | 0,000887 | 0,000000 | 0,000000 | 0,013199 | 0,001249 | 0,001194 | -3,627566 | -19,910093 | -19,640236 |
| **Slc11a2** | 0,167972 | 0,205104 | 0,545038 | 0,196171 | 0,178230 | 0,474967 | 0,024425 | 0,367311 | 0,214432 |
| **Twnk** | 0,264808 | 0,228052 | 0,065079 | 0,480188 | 0,310521 | 0,057371 | -0,589699 | -0,292268 | 0,126636 |
| **Pprc1** | 0,211579 | 0,152672 | 0,042371 | 0,239025 | 0,164445 | 0,011952 | 0,047756 | 0,055281 | 1,753839 |
| **Timm44** | 1,045260 | 1,058413 | 0,628944 | 1,473267 | 1,565895 | 0,646033 | -0,321455 | -0,436429 | -0,007587 |
| **Hdac5** | 0,093465 | 0,062743 | 0,239306 | 0,133193 | 0,062470 | 0,341480 | -0,205962 | 0,157328 | -0,584802 |
| **Max** | 0,545278 | 0,483879 | 2,221689 | 0,698209 | 0,641502 | 3,127712 | -0,111124 | -0,224079 | -0,588663 |
| **Mef2d** | 0,176917 | 0,153353 | 0,383029 | 0,188926 | 0,160334 | 0,439157 | 0,122156 | 0,120992 | -0,181754 |
| **Nr1d2** | 0,359691 | 0,176477 | 0,171831 | 0,258425 | 0,102920 | 0,140902 | 0,733455 | 0,939562 | 0,372006 |
| **Bop1** | 0,324534 | 0,405258 | 0,398432 | 0,319176 | 0,473174 | 0,272599 | 0,242478 | -0,047071 | 0,570025 |
| **Brix1** | 1,088782 | 1,252917 | 0,359117 | 1,330339 | 1,387022 | 0,226979 | -0,083477 | 0,008566 | 0,703258 |
| **Bysl** | 0,192100 | 0,200826 | 0,100615 | 0,199550 | 0,213776 | 0,034383 | 0,141941 | 0,082944 | 1,608121 |
| **Ccdc86** | 0,586440 | 0,580775 | 0,081618 | 0,821472 | 0,791399 | 0,043173 | -0,264446 | -0,290889 | 0,860499 |
| **Ddx1** | 1,491935 | 1,601323 | 1,227196 | 1,650176 | 1,763289 | 1,342371 | 0,059703 | 0,009642 | -0,095234 |

| | | | | | | | | | |
|---|---|---|---|---|---|---|---|---|---|
| **Ddx31** | 0,063106 | 0,061637 | 0,010665 | 0,088861 | 0,079585 | 0,004651 | -0,300899 | -0,151554 | 1,187766 |
| **Dhx37** | 0,095606 | 0,069051 | 0,033048 | 0,093281 | 0,082661 | 0,028004 | 0,215956 | -0,071430 | 0,274961 |
| **Dimt1** | 0,252603 | 0,256481 | 0,028986 | 0,274758 | 0,236680 | 0,031222 | 0,077647 | 0,271806 | 0,053913 |
| **Dkc1** | 0,947010 | 0,997651 | 0,201399 | 1,814779 | 1,629709 | 0,119562 | -0,782569 | -0,575504 | 0,815184 |
| **Exosc2** | 0,327768 | 0,338642 | 0,075000 | 0,372951 | 0,353017 | 0,042026 | 0,025774 | 0,128070 | 0,894059 |
| **Exosc5** | 1,039319 | 0,870564 | 0,466081 | 1,271250 | 1,324574 | 0,509141 | -0,086205 | -0,474579 | -0,085153 |
| **Fbl** | 2,519291 | 2,159544 | 0,512542 | 3,989706 | 3,438841 | 0,322208 | -0,530597 | -0,572538 | 0,736750 |
| **Gnl3l** | 0,353274 | 0,381620 | 0,408428 | 0,402819 | 0,364947 | 0,297939 | 0,057671 | 0,232710 | 0,505753 |
| **Gtpbp4** | 1,410208 | 1,290232 | 0,486537 | 1,809156 | 1,564094 | 0,276832 | -0,158294 | -0,116766 | 0,910197 |
| **Heatr1** | 0,234528 | 0,229555 | 0,087421 | 0,244935 | 0,244405 | 0,021247 | 0,167494 | 0,107005 | 2,190218 |
| **Imp4** | 0,849255 | 0,842353 | 0,342653 | 1,126425 | 1,223793 | 0,301076 | -0,210811 | -0,386070 | 0,216382 |
| **Ltv1** | 0,876299 | 0,834838 | 0,504844 | 0,739697 | 0,690809 | 0,203828 | 0,500884 | 0,476723 | 1,322273 |
| **Mak16** | 0,818796 | 0,749502 | 0,409568 | 1,240875 | 1,080616 | 0,237300 | -0,385464 | -0,382623 | 0,945204 |
| **Mdn1** | 0,310328 | 0,200309 | 0,067677 | 0,369930 | 0,243573 | 0,023588 | -0,064019 | -0,140706 | 1,557617 |
| **Mrto4** | 1,008828 | 1,055499 | 0,153375 | 1,500934 | 1,502623 | 0,094360 | -0,389054 | -0,358913 | 0,707488 |
| **Naf1** | 0,268687 | 0,188301 | 0,032588 | 0,426927 | 0,245390 | 0,020416 | -0,436975 | -0,225498 | 0,671992 |
| **Nhp2** | 2,565105 | 2,947262 | 0,442321 | 3,924238 | 4,484311 | 0,243629 | -0,505989 | -0,522405 | 1,016417 |
| **Nle1** | 0,135206 | 0,119286 | 0,016088 | 0,309518 | 0,224193 | 0,007503 | -0,947794 | -0,730890 | 0,764166 |
| **Nob1** | 0,432084 | 0,380035 | 0,153735 | 0,718195 | 0,536029 | 0,102764 | -0,515783 | -0,349394 | 0,858628 |
| **Noc2l** | 0,745068 | 0,572124 | 0,183216 | 1,110469 | 0,820605 | 0,085230 | -0,377623 | -0,355963 | 1,256990 |
| **Nol6** | 0,148747 | 0,147287 | 0,061981 | 0,168423 | 0,163271 | 0,034694 | 0,065234 | 0,011038 | 0,845518 |
| **Nop2** | 0,288885 | 0,270265 | 0,114395 | 0,454468 | 0,339829 | 0,146916 | -0,436721 | -0,150828 | -0,293111 |
| **Nop56** | 1,675124 | 1,559836 | 0,209715 | 2,241613 | 1,863932 | 0,137724 | -0,243641 | -0,071350 | 0,830048 |
| **Nop58** | 2,667917 | 2,619704 | 0,714237 | 3,397794 | 2,922302 | 0,430382 | -0,185775 | -0,030102 | 0,780341 |
| **Pdcd11** | 0,512393 | 0,446010 | 0,348518 | 0,596454 | 0,491422 | 0,245874 | 0,005758 | -0,005553 | 0,554522 |
| **Pes1** | 0,517615 | 0,457871 | 0,257928 | 0,747096 | 0,655739 | 0,225196 | -0,312381 | -0,370449 | 0,348683 |
| **Ppan** | 0,429579 | 0,336848 | 0,101189 | 0,736732 | 0,595855 | 0,037682 | -0,585708 | -0,689420 | 1,325830 |
| **Rcl1** | 0,379194 | 0,303779 | 0,053019 | 0,747279 | 0,541539 | 0,079525 | -0,759720 | -0,674710 | -0,426409 |
| **Riok1** | 0,404608 | 0,441033 | 0,342811 | 0,455594 | 0,455968 | 0,312882 | 0,011956 | 0,110850 | 0,103332 |
| **Rpl10a** | 31,213152 | 24,189297 | 8,742778 | 75,636658 | 56,922882 | 13,927016 | -1,316626 | -1,263692 | -0,455356 |
| **Rpl13** | 66,824997 | 55,756733 | 17,128996 | 146,891830 | 120,546616 | 24,867874 | -1,164488 | -1,138248 | -0,273564 |
| **Rpl17** | 38,375824 | 30,701231 | 9,441228 | 75,459991 | 57,538658 | 13,396972 | -1,018258 | -0,947961 | -0,447894 |
| **Rpl22l1** | 8,492740 | 7,568616 | 2,189115 | 22,438156 | 19,590851 | 4,908749 | -1,554921 | -1,534636 | -1,121169 |
| **Rpl23** | 51,896084 | 41,995903 | 15,271737 | 119,908829 | 92,486038 | 24,003019 | -1,244260 | -1,166860 | -0,542232 |
| **Rpl23a** | 19,546684 | 17,355835 | 5,129654 | 33,937366 | 28,620806 | 7,097049 | -0,805226 | -0,748723 | -0,320740 |
| **Rpl24** | 34,371613 | 29,746412 | 13,370777 | 59,995426 | 48,487450 | 16,084803 | -0,827721 | -0,732055 | -0,212274 |
| **Rpl29** | 24,485271 | 22,942179 | 10,933121 | 47,862118 | 43,654827 | 12,959869 | -1,020206 | -0,988369 | -0,164385 |
| **Rpl32** | 51,897465 | 42,786854 | 12,025021 | 120,553963 | 96,766113 | 19,600319 | -1,263514 | -1,222538 | -0,496704 |
| **Rpl34** | 37,011826 | 31,039314 | 11,930344 | 73,901085 | 59,895641 | 16,474888 | -1,056352 | -1,002623 | -0,395849 |
| **Rpl35a** | 43,423763 | 37,316921 | 13,268756 | 85,083900 | 69,494820 | 19,190691 | -1,005143 | -0,930435 | -0,476792 |
| **Rpl37a** | 42,883255 | 34,636299 | 14,546797 | 86,687859 | 67,679352 | 21,414536 | -1,052943 | -1,000989 | -0,507493 |
| **Rpl7l1** | 1,277392 | 1,390670 | 0,748299 | 1,394244 | 1,576376 | 0,684006 | 0,082915 | -0,024252 | 0,188414 |
| **Rps12** | 55,627285 | 45,506676 | 15,874035 | 131,994766 | 105,858223 | 27,110006 | -1,282876 | -1,254883 | -0,654995 |
| **Rps2** | 54,679565 | 46,267494 | 11,854826 | 116,610870 | 98,889793 | 14,984224 | -1,185393 | -1,160541 | -0,227235 |

| | | | | | | | | | |
|---|---|---|---|---|---|---|---|---|---|
| **Rps20** | 56,708698 | 44,126163 | 14,334633 | 139,438782 | 109,986275 | 28,349466 | -1,345761 | -1,353810 | -0,970841 |
| **Rps21** | 38,158276 | 31,493048 | 9,437173 | 80,973457 | 62,054886 | 14,504363 | -1,124507 | -1,003472 | -0,475617 |
| **Rps24** | 57,679974 | 47,250748 | 15,695683 | 128,701752 | 101,786766 | 27,858583 | -1,187975 | -1,138055 | -0,585883 |
| **Rps25** | 24,226755 | 20,787252 | 9,036367 | 41,204948 | 34,160450 | 9,299698 | -0,778700 | -0,735333 | 0,043229 |
| **Rps3** | 37,175388 | 31,155014 | 10,241975 | 78,925255 | 60,737305 | 14,297837 | -1,143634 | -0,998573 | -0,470278 |
| **Rps7** | 39,994545 | 32,359329 | 9,532992 | 91,553741 | 69,749367 | 14,935725 | -1,243786 | -1,150685 | -0,496654 |
| **Rrp1** | 2,203742 | 2,176899 | 1,506586 | 2,408706 | 2,260630 | 1,190868 | 0,063737 | 0,071728 | 0,405919 |
| **Rrp1b** | 0,273490 | 0,179014 | 0,014311 | 0,498788 | 0,377054 | 0,015237 | -0,691960 | -0,892287 | 0,129316 |
| **Rrs1** | 0,654848 | 0,564086 | 0,088863 | 1,062147 | 0,755727 | 0,063914 | -0,490754 | -0,240977 | 0,719447 |
| **Rsl1d1** | 1,928784 | 1,694692 | 0,555396 | 2,730576 | 2,340346 | 0,343709 | -0,336505 | -0,329733 | 0,674209 |
| **Surf6** | 0,207296 | 0,183382 | 0,088726 | 0,252472 | 0,242444 | 0,078478 | -0,062869 | -0,243329 | 0,253178 |
| **Tex10** | 0,321189 | 0,321966 | 0,169687 | 0,413044 | 0,354200 | 0,079256 | -0,151002 | 0,028954 | 1,044391 |
| **Utp14a** | 0,747413 | 0,644968 | 0,183082 | 1,589280 | 1,038685 | 0,144550 | -0,911777 | -0,529412 | 0,503180 |
| **Utp18** | 0,664818 | 0,588339 | 0,237588 | 0,778489 | 0,675183 | 0,146626 | 0,008067 | -0,017010 | 0,737039 |
| **Utp20** | 0,442610 | 0,321583 | 0,121012 | 0,422602 | 0,308085 | 0,046260 | 0,310982 | 0,211262 | 1,478337 |
| **Wdr3** | 0,310553 | 0,232361 | 0,163520 | 0,345501 | 0,294444 | 0,069514 | 0,051720 | -0,132857 | 1,238174 |
| **Wdr43** | 1,287279 | 1,079060 | 0,317207 | 1,874436 | 1,470720 | 0,201040 | -0,351593 | -0,304491 | 0,751568 |
| **Wdr74** | 0,596106 | 0,624976 | 0,288112 | 0,855043 | 0,898362 | 0,186092 | -0,310650 | -0,374557 | 0,726955 |
| **Wdr75** | 0,559959 | 0,523820 | 0,088898 | 0,689782 | 0,618118 | 0,047510 | -0,072966 | -0,047528 | 0,912461 |
| **Dazap1** | 1,533183 | 1,894695 | 1,049433 | 1,633926 | 1,997468 | 0,868297 | 0,121971 | 0,074972 | 0,364050 |
| **Ddx10** | 0,661599 | 0,610368 | 0,374211 | 0,701981 | 0,567071 | 0,147741 | 0,121890 | 0,259821 | 1,202284 |
| **Ddx18** | 1,544915 | 1,449549 | 0,506867 | 1,728706 | 1,624163 | 0,317049 | 0,041130 | 0,001440 | 0,691975 |
| **Ddx55** | 0,367743 | 0,349102 | 0,246735 | 0,425589 | 0,343760 | 0,152572 | -0,035887 | 0,200832 | 0,574174 |
| **Dus3l** | 0,181331 | 0,174957 | 0,146626 | 0,278328 | 0,263415 | 0,137692 | -0,429293 | -0,392227 | 0,117935 |
| **Eif3b** | 1,452830 | 1,499334 | 1,125543 | 1,503609 | 1,562323 | 0,799342 | 0,149297 | 0,100434 | 0,558252 |
| **Elp2** | 0,289079 | 0,266378 | 0,216782 | 0,336225 | 0,312755 | 0,128543 | 0,027065 | -0,054250 | 0,794748 |
| **Mettl1** | 0,922343 | 0,634250 | 0,105678 | 1,335371 | 1,007837 | 0,062804 | -0,341441 | -0,550050 | 0,978660 |
| **Nifk** | 0,780250 | 0,727465 | 0,186973 | 1,249271 | 0,973682 | 0,160237 | -0,467247 | -0,280323 | 0,305612 |
| **Polr1b** | 0,138233 | 0,101226 | 0,030536 | 0,205885 | 0,177934 | 0,011176 | -0,378514 | -0,568305 | 1,380364 |
| **Polr1c** | 0,609681 | 0,536660 | 0,347034 | 0,764255 | 0,671120 | 0,453737 | -0,108007 | -0,151139 | -0,429901 |
| **Pus1** | 0,728466 | 0,610682 | 0,277539 | 1,086530 | 0,946447 | 0,258785 | -0,364626 | -0,488368 | 0,212548 |
| **Trmt61a** | 0,179167 | 0,162816 | 0,038319 | 0,237681 | 0,221631 | 0,022577 | -0,173929 | -0,255381 | 0,862188 |
| **Btf3** | 15,767461 | 14,420588 | 8,160940 | 21,678091 | 19,099142 | 8,991222 | -0,490400 | -0,425841 | -0,148916 |
| **Spout1** | 0,350403 | 0,273355 | 0,136297 | 0,408611 | 0,332083 | 0,101221 | 0,016725 | -0,141709 | 0,527294 |
| **Nop53** | 1,494078 | 1,275150 | 1,084806 | 2,377143 | 2,008241 | 1,625733 | -0,560551 | -0,543236 | -0,668717 |
| **Gpatch4** | 0,563377 | 0,550516 | 0,100479 | 1,000082 | 0,940982 | 0,049510 | -0,645304 | -0,619253 | 1,073295 |
| **Grwd1** | 0,255422 | 0,268870 | 0,033825 | 0,462362 | 0,388580 | 0,012121 | -0,608958 | -0,342280 | 1,541721 |
| **Pum3** | 0,860413 | 0,704395 | 0,410604 | 1,163804 | 0,945113 | 0,205926 | -0,227357 | -0,260600 | 0,948668 |
| **Klhdc4** | 0,334072 | 0,454299 | 0,306184 | 0,342602 | 0,451862 | 0,227085 | 0,186164 | 0,179954 | 0,422369 |
| **Noc3l** | 0,374246 | 0,310468 | 0,070164 | 0,446381 | 0,375119 | 0,037376 | -0,037067 | -0,088928 | 0,903004 |
| **Rrp12** | 0,098197 | 0,084877 | 0,046701 | 0,098528 | 0,078950 | 0,016905 | 0,186020 | 0,236458 | 1,385865 |
| **Tma16** | 0,456886 | 0,324052 | 0,072695 | 0,614969 | 0,450893 | 0,039715 | -0,166784 | -0,308024 | 0,774317 |
| **Urb1** | 0,092115 | 0,074744 | 0,016110 | 0,091798 | 0,054578 | 0,016778 | 0,247811 | 0,628314 | 0,211838 |
| **Polr1f** | 0,000000 | 0,000000 | 0,000000 | 0,000000 | 0,000000 | 0,000000 | 0,000000 | 0,000000 | 0,000000 |

2 days post irradiation (2dpi)

| Genotype | WT (2dpi) | | | $Myc^{\Delta 2\text{-}540/\Delta 2\text{-}540}$ (2dpi) | | | Fold change (log2) WT (2dpi) *vs* $Myc^{\Delta 2\text{-}540/\Delta 2\text{-}540}$ (2dpi) | | |
|---|---|---|---|---|---|---|---|---|---|
| Gene / Cluster | SC | TA | EC | SC | TA | EC | SC | TA | EC |
| **Atic** | 2,351068 | 2,901829 | 0,098874 | 0,928649 | 0,881183 | 0,048115 | 1,613715 | 1,995323 | 1,111949 |
| **Ctps** | 0,727709 | 0,871389 | 0,012937 | 0,263640 | 0,283589 | 0,003500 | 1,746151 | 1,835980 | 2,055819 |
| **Ctps2** | 0,137893 | 0,112348 | 0,055180 | 0,227268 | 0,167519 | 0,062204 | -0,557998 | -0,367570 | -0,037767 |
| **Nme1** | 16,894339 | 19,010582 | 1,803144 | 8,589489 | 9,219363 | 4,266795 | 0,976450 | 1,136604 | -1,478295 |
| **Nme2** | 30,735613 | 32,834606 | 5,474013 | 16,380322 | 16,483839 | 14,729856 | 0,877973 | 1,040390 | -1,833149 |
| **Ppat** | 1,164496 | 1,167984 | 0,003881 | 0,311504 | 0,190704 | 0,000000 | 2,248294 | 2,887820 | 21,159414 |
| **Ahcy** | 0,273737 | 0,357180 | 0,063840 | 0,182316 | 0,214901 | 0,085320 | 0,787312 | 0,945089 | -0,417072 |
| **Shmt2** | 0,718720 | 0,942808 | 0,020820 | 0,110848 | 0,199622 | 0,011724 | 2,941833 | 2,574511 | 0,907693 |
| **Aldh7a1** | 0,367763 | 0,342216 | 0,168137 | 0,259146 | 0,225856 | 0,094006 | 0,706983 | 0,868519 | 0,902015 |
| **Glo1** | 3,343735 | 3,236774 | 2,545915 | 3,440486 | 3,407493 | 4,488729 | 0,060336 | 0,062822 | -1,044469 |
| **Pck1** | 0,014435 | 0,000788 | 0,651113 | 0,015769 | 0,002874 | 0,437791 | 0,018230 | -1,660518 | 0,258261 |
| **Srm** | 1,867309 | 2,617611 | 0,035900 | 0,184340 | 0,209350 | 0,023740 | 3,621039 | 3,967779 | 0,370813 |
| **Acp2** | 0,116511 | 0,085999 | 0,323286 | 0,214764 | 0,122006 | 0,410455 | -0,656714 | -0,292952 | -0,411028 |
| **Dph5** | 0,478788 | 0,482557 | 0,066876 | 0,275797 | 0,286050 | 0,104208 | 1,015802 | 1,040585 | -0,437549 |
| **Hexa** | 0,887636 | 0,766941 | 2,015169 | 1,621659 | 1,570404 | 2,446416 | -0,796075 | -0,877874 | -0,348227 |
| **Dhrs3** | 0,150014 | 0,072727 | 0,346925 | 0,297273 | 0,237629 | 0,534405 | -0,936339 | -1,609391 | -0,668943 |
| **Slc2a6** | 0,000000 | 0,000362 | 0,002075 | 0,005862 | 0,002830 | 0,000000 | -21,870047 | -2,608100 | 20,227404 |
| **Hspe1** | 17,033276 | 19,972420 | 0,969065 | 10,714465 | 10,251244 | 2,450710 | 0,662821 | 1,045793 | -1,440134 |
| **Prmt3** | 0,348778 | 0,452988 | 0,010173 | 0,124245 | 0,147574 | 0,010497 | 1,705702 | 1,804648 | 0,103534 |
| **Psme2** | 3,216036 | 2,839527 | 6,446042 | 5,407202 | 5,172930 | 8,796909 | -0,746509 | -0,824171 | -0,697071 |
| **Rab11fip1** | 0,748319 | 0,550101 | 2,392028 | 0,894474 | 0,667818 | 1,426970 | -0,215708 | -0,119929 | 0,600440 |
| **Uevld** | 0,105475 | 0,114425 | 0,128823 | 0,090765 | 0,254483 | 0,147639 | 0,341253 | -0,930692 | -0,200451 |
| **Usp2** | 0,115528 | 0,078602 | 0,597028 | 0,221221 | 0,180767 | 0,986020 | -0,721238 | -0,932765 | -0,760057 |
| **Tnfaip8l1** | 0,316550 | 0,393870 | 0,066916 | 0,513178 | 0,684168 | 0,032330 | -0,498721 | -0,610124 | 1,034078 |
| **Gulp1** | 0,000000 | 0,000000 | 0,000000 | 0,000000 | 0,000000 | 0,000000 | 0,000000 | 0,000000 | 0,000000 |
| **Slc11a2** | 0,170995 | 0,165727 | 0,263998 | 0,264587 | 0,184088 | 0,595004 | -0,496235 | 0,079526 | -1,092962 |
| **Twnk** | 0,312355 | 0,363280 | 0,027968 | 0,173070 | 0,165780 | 0,069673 | 1,121268 | 1,400893 | -1,296058 |
| **Pprc1** | 0,307231 | 0,395991 | 0,009289 | 0,178135 | 0,161961 | 0,037072 | 1,049821 | 1,550878 | -1,929907 |
| **Timm44** | 1,300936 | 1,540437 | 0,373424 | 0,909021 | 0,882679 | 0,593146 | 0,732780 | 1,030456 | -0,652079 |
| **Hdac5** | 0,023516 | 0,027870 | 0,137392 | 0,100453 | 0,042238 | 0,385387 | -1,826494 | -0,452006 | -1,629799 |
| **Max** | 0,667796 | 0,625252 | 3,351505 | 0,464046 | 0,364189 | 2,082839 | 0,699148 | 1,031322 | 0,694981 |
| **Mef2d** | 0,228017 | 0,189531 | 0,461184 | 0,318175 | 0,195303 | 0,424823 | -0,261058 | 0,130314 | 0,214033 |
| **Nr1d2** | 0,196854 | 0,141318 | 0,342424 | 0,275202 | 0,230515 | 0,245846 | -0,406369 | -0,476386 | 0,344053 |
| **Bop1** | 0,761877 | 0,892882 | 0,253038 | 0,397162 | 0,514128 | 0,343658 | 1,114211 | 1,049273 | -0,422998 |
| **Brix1** | 2,280455 | 2,784763 | 0,263844 | 1,273848 | 1,513285 | 0,369750 | 1,001295 | 1,073310 | -0,507429 |
| **Bysl** | 0,581908 | 0,741740 | 0,039280 | 0,214048 | 0,236733 | 0,062339 | 1,679172 | 1,905531 | -0,885915 |
| **Ccdc86** | 1,821635 | 2,024815 | 0,049723 | 0,707024 | 0,724915 | 0,067860 | 1,594521 | 1,768385 | -0,350404 |
| **Ddx1** | 1,826090 | 1,965890 | 1,271748 | 1,280561 | 1,424135 | 1,029241 | 0,709339 | 0,663813 | 0,302906 |
| **Ddx31** | 0,115626 | 0,181199 | 0,002811 | 0,054279 | 0,053515 | 0,021056 | 1,326570 | 1,975210 | -2,431216 |
| **Dhx37** | 0,165073 | 0,145245 | 0,043693 | 0,067680 | 0,082567 | 0,022260 | 1,523853 | 0,982656 | 0,877103 |
| **Dimt1** | 0,470523 | 0,605875 | 0,016481 | 0,228006 | 0,292389 | 0,019891 | 1,256224 | 1,305371 | -0,201109 |

| | | | | | | | | | |
|---|---|---|---|---|---|---|---|---|---|
| **Dkc1** | 2,538009 | 3,227853 | 0,052161 | 0,718559 | 0,854304 | 0,059959 | 2,123837 | 2,250884 | -0,087061 |
| **Exosc2** | 0,456438 | 0,518719 | 0,013180 | 0,197036 | 0,216333 | 0,020739 | 1,525054 | 1,517160 | -0,538571 |
| **Exosc5** | 1,418237 | 1,613709 | 0,375112 | 0,856946 | 0,917817 | 0,475946 | 0,963274 | 1,051548 | -0,284143 |
| **Fbl** | 4,753687 | 5,446901 | 0,253112 | 1,874044 | 2,010055 | 0,226146 | 1,508814 | 1,665698 | 0,119452 |
| **Gnl3l** | 0,511184 | 0,490356 | 0,340659 | 0,284966 | 0,253490 | 0,231435 | 1,020088 | 1,149684 | 0,484885 |
| **Gtpbp4** | 2,534945 | 3,086287 | 0,235875 | 1,000024 | 1,205784 | 0,360645 | 1,571763 | 1,614933 | -0,660566 |
| **Heatr1** | 0,532697 | 0,649278 | 0,057904 | 0,191496 | 0,194140 | 0,030748 | 1,647781 | 1,962914 | 0,572419 |
| **Imp4** | 1,448952 | 1,824067 | 0,304937 | 0,836719 | 0,851144 | 0,419997 | 1,055887 | 1,319221 | -0,368443 |
| **Ltv1** | 0,948945 | 1,170018 | 0,370850 | 0,998278 | 1,008556 | 0,481807 | 0,154954 | 0,415712 | -0,321542 |
| **Mak16** | 2,094352 | 2,345347 | 0,208816 | 0,742348 | 0,784944 | 0,276902 | 1,733683 | 1,859900 | -0,421488 |
| **Mdn1** | 0,710353 | 0,904390 | 0,025954 | 0,328684 | 0,195414 | 0,015877 | 1,363202 | 2,446526 | 0,576310 |
| **Mrto4** | 3,156299 | 3,942912 | 0,119126 | 0,963478 | 1,243238 | 0,167021 | 1,972034 | 1,933579 | -0,451109 |
| **Naf1** | 0,693234 | 0,919542 | 0,042960 | 0,202606 | 0,209806 | 0,019683 | 1,973190 | 2,368491 | 1,044356 |
| **Nhp2** | 7,931756 | 10,110726 | 0,319690 | 2,799663 | 3,791767 | 0,469794 | 1,688478 | 1,565650 | -0,612167 |
| **Nle1** | 0,415242 | 0,553820 | 0,002101 | 0,082437 | 0,127474 | 0,006200 | 2,541542 | 2,393388 | -1,541621 |
| **Nob1** | 0,653807 | 0,815665 | 0,053554 | 0,298834 | 0,312951 | 0,065543 | 1,416656 | 1,677308 | -0,210031 |
| **Noc2l** | 1,409950 | 1,709387 | 0,058704 | 0,580494 | 0,568943 | 0,074728 | 1,503643 | 1,866911 | -0,208045 |
| **Nol6** | 0,213194 | 0,222368 | 0,018847 | 0,086767 | 0,077652 | 0,040896 | 1,480317 | 1,816340 | -0,880878 |
| **Nop2** | 0,745600 | 0,919550 | 0,124730 | 0,277040 | 0,258922 | 0,180536 | 1,709546 | 2,105729 | -0,482121 |
| **Nop56** | 3,941887 | 4,704150 | 0,159267 | 1,431268 | 1,590253 | 0,127258 | 1,773975 | 1,858115 | 0,253367 |
| **Nop58** | 4,106544 | 5,731502 | 0,585695 | 1,836580 | 2,274814 | 0,457198 | 1,403983 | 1,580953 | 0,316917 |
| **Pdcd11** | 0,946214 | 0,951587 | 0,134316 | 0,437246 | 0,390152 | 0,364415 | 1,291277 | 1,529233 | -1,389242 |
| **Pes1** | 0,903326 | 1,019976 | 0,137326 | 0,355344 | 0,370325 | 0,169781 | 1,596482 | 1,690414 | -0,303654 |
| **Ppan** | 1,150624 | 1,505602 | 0,028123 | 0,406073 | 0,398762 | 0,060041 | 1,846371 | 2,228244 | -0,899535 |
| **Rcl1** | 0,732694 | 0,899572 | 0,034048 | 0,275878 | 0,256344 | 0,047730 | 1,666497 | 2,116250 | -0,274379 |
| **Riok1** | 0,599814 | 0,659391 | 0,274576 | 0,392661 | 0,455401 | 0,243384 | 0,763039 | 0,752944 | 0,136277 |
| **Rpl10a** | 46,493034 | 46,026306 | 8,229234 | 26,969015 | 24,047476 | 7,611582 | 0,752739 | 0,976107 | -0,103093 |
| **Rpl13** | 120,383591 | 121,256844 | 14,632399 | 70,619690 | 67,456856 | 14,035484 | 0,735221 | 0,879328 | -0,043698 |
| **Rpl17** | 53,250542 | 53,205654 | 10,888444 | 34,191975 | 32,222244 | 10,623349 | 0,609422 | 0,763892 | -0,057322 |
| **Rpl22l1** | 7,505990 | 9,726561 | 2,335294 | 4,840593 | 5,479592 | 2,899729 | 0,593562 | 0,988750 | -0,385443 |
| **Rpl23** | 63,100922 | 61,390884 | 12,492678 | 38,261265 | 35,282101 | 12,073855 | 0,689823 | 0,833628 | -0,040596 |
| **Rpl23a** | 28,092649 | 29,391706 | 4,487409 | 11,812597 | 11,504436 | 3,660062 | 1,317403 | 1,465058 | 0,296295 |
| **Rpl24** | 37,951675 | 39,107586 | 10,326513 | 27,219469 | 27,720200 | 12,210615 | 0,471864 | 0,525802 | -0,339672 |
| **Rpl29** | 40,315746 | 41,954468 | 8,915174 | 20,918774 | 22,184393 | 9,537887 | 0,915374 | 0,969307 | -0,220826 |
| **Rpl32** | 65,433548 | 66,118370 | 9,263281 | 35,303020 | 36,247387 | 8,407339 | 0,878677 | 0,909226 | 0,063111 |
| **Rpl34** | 31,518950 | 31,614334 | 7,171304 | 17,894983 | 17,555555 | 7,929123 | 0,854645 | 0,914441 | -0,208632 |
| **Rpl35a** | 31,979326 | 33,046494 | 6,166638 | 21,226542 | 20,811577 | 7,937500 | 0,593807 | 0,723104 | -0,432493 |
| **Rpl37a** | 27,884809 | 26,541571 | 5,709682 | 17,746374 | 17,037121 | 5,977150 | 0,678054 | 0,699057 | -0,106541 |
| **Rpl7l1** | 1,967897 | 2,643416 | 0,633201 | 1,295575 | 1,480256 | 0,991997 | 0,785783 | 1,055624 | -0,676779 |
| **Rps12** | 80,831703 | 81,993416 | 12,792134 | 31,413586 | 30,644499 | 10,437498 | 1,352059 | 1,453934 | 0,273560 |
| **Rps2** | 116,563965 | 130,494904 | 10,735313 | 57,704498 | 59,550175 | 13,422683 | 0,977189 | 1,163689 | -0,457973 |
| **Rps20** | 64,411774 | 64,608765 | 20,786400 | 40,626278 | 39,035168 | 13,668818 | 0,636674 | 0,752068 | 0,567659 |
| **Rps21** | 24,422890 | 24,318689 | 3,137450 | 14,106364 | 13,067758 | 3,488823 | 0,837890 | 0,970430 | -0,213207 |
| **Rps24** | 60,428242 | 57,931728 | 9,761638 | 38,523659 | 36,583221 | 8,978566 | 0,640111 | 0,686960 | -0,115855 |

| | | | | | | | | | |
|---|---|---|---|---|---|---|---|---|---|
| **Rps25** | 23,784372 | 23,938208 | 5,332410 | 11,350204 | 10,949293 | 5,160457 | 1,125978 | 1,217259 | -0,001478 |
| **Rps3** | 70,101837 | 71,043396 | 12,781179 | 41,896244 | 42,636612 | 12,365615 | 0,702829 | 0,748221 | -0,034726 |
| **Rps7** | 64,854996 | 64,965118 | 10,399237 | 41,033684 | 39,350677 | 10,075089 | 0,605872 | 0,734177 | -0,119869 |
| **Rrp1** | 2,758522 | 3,193993 | 1,283081 | 2,645614 | 2,686640 | 1,617069 | 0,194979 | 0,402405 | -0,305919 |
| **Rrp1b** | 1,267145 | 1,552789 | 0,010941 | 0,206218 | 0,188890 | 0,009961 | 2,861465 | 3,346333 | 0,438562 |
| **Rrs1** | 1,493464 | 1,710270 | 0,106881 | 0,578670 | 0,577904 | 0,043890 | 1,630627 | 1,832471 | 1,201070 |
| **Rsl1d1** | 4,565586 | 5,316899 | 0,261298 | 1,617649 | 1,640147 | 0,427066 | 1,693577 | 1,960167 | -0,675128 |
| **Surf6** | 0,312500 | 0,314161 | 0,042344 | 0,130975 | 0,123933 | 0,060676 | 1,443212 | 1,632022 | -0,370110 |
| **Tex10** | 0,467222 | 0,566851 | 0,061428 | 0,242426 | 0,274285 | 0,078524 | 1,168466 | 1,279215 | -0,247706 |
| **Utp14a** | 1,508083 | 1,744274 | 0,106166 | 0,656996 | 0,640177 | 0,105194 | 1,414178 | 1,695817 | -0,054936 |
| **Utp18** | 1,015463 | 1,338896 | 0,194913 | 0,446316 | 0,591721 | 0,217588 | 1,435551 | 1,424270 | -0,258192 |
| **Utp20** | 0,957203 | 1,069432 | 0,030559 | 0,597629 | 0,480248 | 0,014136 | 0,882922 | 1,407136 | 1,045659 |
| **Wdr3** | 0,572880 | 0,691565 | 0,045738 | 0,192276 | 0,216289 | 0,044246 | 1,831783 | 1,954558 | 0,192027 |
| **Wdr43** | 2,192452 | 2,684169 | 0,121472 | 0,714570 | 0,808057 | 0,176886 | 1,856799 | 1,995254 | -0,625903 |
| **Wdr74** | 1,338244 | 1,877745 | 0,138916 | 0,685862 | 0,820135 | 0,313273 | 1,134476 | 1,448820 | -1,135063 |
| **Wdr75** | 0,947070 | 1,251544 | 0,026312 | 0,396328 | 0,472814 | 0,037517 | 1,516114 | 1,700228 | -0,414588 |
| **Dazap1** | 1,588313 | 1,882696 | 0,516223 | 1,261632 | 1,677746 | 0,708721 | 0,462647 | 0,361328 | -0,473738 |
| **Ddx10** | 0,918101 | 0,974357 | 0,127693 | 0,500570 | 0,562008 | 0,139670 | 1,045969 | 1,021450 | -0,134725 |
| **Ddx18** | 1,544271 | 1,946361 | 0,165162 | 0,767298 | 0,820602 | 0,240368 | 1,254654 | 1,556858 | -0,419101 |
| **Ddx55** | 0,708794 | 0,540880 | 0,187886 | 0,344287 | 0,353997 | 0,200436 | 1,299093 | 0,825064 | -0,073369 |
| **Dus3l** | 0,226635 | 0,267753 | 0,093715 | 0,170211 | 0,178453 | 0,117423 | 0,587587 | 0,844633 | -0,248619 |
| **Eif3b** | 2,768667 | 3,431437 | 0,712211 | 1,601342 | 1,542133 | 0,932359 | 0,977465 | 1,353024 | -0,415422 |
| **Elp2** | 0,277251 | 0,354534 | 0,132891 | 0,233655 | 0,214983 | 0,118956 | 0,509671 | 0,952775 | 0,161740 |
| **Mettl1** | 1,298998 | 1,935464 | 0,042910 | 0,647702 | 0,670431 | 0,054685 | 1,283359 | 1,871310 | -0,421119 |
| **Nifk** | 1,476723 | 1,726248 | 0,089607 | 0,647769 | 0,663500 | 0,145975 | 1,468773 | 1,629283 | -0,616144 |
| **Polr1b** | 0,301479 | 0,426187 | 0,007638 | 0,108430 | 0,101430 | 0,010081 | 1,773044 | 2,351280 | -0,208234 |
| **Polr1c** | 1,097043 | 1,171173 | 0,454244 | 0,587082 | 0,603309 | 0,273829 | 1,115353 | 1,235181 | 0,756530 |
| **Pus1** | 1,258256 | 1,471615 | 0,247029 | 0,774253 | 0,749350 | 0,386025 | 0,857160 | 1,213052 | -0,708243 |
| **Trmt61a** | 0,302625 | 0,508266 | 0,010611 | 0,140619 | 0,189891 | 0,040471 | 1,338217 | 1,722147 | -2,143686 |
| **Btf3** | 21,308796 | 23,132681 | 10,682523 | 17,521801 | 18,080755 | 10,577289 | 0,271303 | 0,389300 | -0,136867 |
| **Spout1** | 0,578723 | 0,670606 | 0,142379 | 0,285471 | 0,269425 | 0,165959 | 1,264506 | 1,581344 | -0,182940 |
| **Nop53** | 1,916817 | 1,961630 | 1,286473 | 1,230575 | 1,224006 | 1,128526 | 0,903141 | 0,898348 | 0,258165 |
| **Gpatch4** | 1,639213 | 2,120152 | 0,082337 | 0,437585 | 0,559209 | 0,061537 | 2,168194 | 2,253076 | 0,283919 |
| **Grwd1** | 0,654502 | 0,898583 | 0,024090 | 0,222329 | 0,345546 | 0,021524 | 1,824546 | 1,666727 | 0,316367 |
| **Pum3** | 1,376032 | 1,534610 | 0,200248 | 0,622216 | 0,621246 | 0,231458 | 1,363131 | 1,550370 | -0,248942 |
| **Klhdc4** | 0,660566 | 0,783213 | 0,277010 | 0,419391 | 0,544680 | 0,271236 | 0,889651 | 0,728592 | 0,042334 |
| **Noc3l** | 0,519558 | 0,779257 | 0,046734 | 0,276420 | 0,292035 | 0,059662 | 1,156101 | 1,631679 | -0,243512 |
| **Rrp12** | 0,242773 | 0,364208 | 0,030661 | 0,111544 | 0,116769 | 0,043235 | 1,365673 | 1,914584 | -0,578866 |
| **Tma16** | 0,807413 | 1,037726 | 0,023202 | 0,344707 | 0,340566 | 0,039676 | 1,508915 | 1,883221 | -0,673782 |
| **Urb1** | 0,180926 | 0,204682 | 0,006115 | 0,070529 | 0,065846 | 0,010815 | 1,500847 | 1,847223 | -0,669648 |
| **Polr1f** | 0,000000 | 0,000000 | 0,000000 | 0,000000 | 0,000000 | 0,000000 | 0,000000 | 0,000000 | 0,000000 |

**b:** Mean expression (Normalised read counts) of Myc family members within SC, TA and EC clusters

Non-irradiated

| Genotype | $Myc^{\Delta 2\text{-}540/\Delta 2\text{-}540}$ | | | WT | | | Fold change (log2) $Myc^{\Delta 2\text{-}540/\Delta 2\text{-}540}$ *vs* WT | | |
|---|---|---|---|---|---|---|---|---|---|
| Gene / Cluster | SC | TA | EC | SC | TA | EC | SC | TA | EC |
| **Myc** | 0,027570 | 0,006726 | 0,000000 | 0,982967 | 0,469614 | 0,021379 | -5,104638 | -6,019047 | -23,361252 |
| **Mycl** | 0,726756 | 0,413762 | 0,071377 | 0,418907 | 0,269462 | 0,050007 | 1,023310 | 0,776771 | 0,503953 |
| **Mycn** | 0,070654 | 0,061945 | 0,006757 | 0,039120 | 0,047735 | 0,002313 | 1,079653 | 0,580262 | 1,469525 |

2 days post irradiation (2dpi)

| Genotype | WT (2dpi) | | | $Myc^{\Delta 2\text{-}540/\Delta 2\text{-}540}$ (2dpi) | | | Fold change (log2) WT (2dpi) *vs* $Myc^{\Delta 2\text{-}540/\Delta 2\text{-}540}$ (2dpi) | | |
|---|---|---|---|---|---|---|---|---|---|
| Gene / Cluster | SC | TA | EC | SC | TA | EC | SC | TA | EC |
| **Myc** | 1,375095 | 1,142745 | 0,027782 | 0,007331 | 0,000829 | 0,000000 | 7,978114 | 10,402727 | 23,852833 |
| **Mycl** | 0,140641 | 0,144756 | 0,005229 | 0,346832 | 0,271238 | 0,014483 | -1,144095 | -0,650227 | -1,290220 |
| **Mycn** | 0,008906 | 0,002044 | 0,000520 | 0,025866 | 0,026860 | 0,000912 | -1,288067 | -3,447741 | -0,620870 |
